# Autotransporters drive biofilm formation and auto-aggregation in the diderm Firmicute *Veillonella parvula*

**DOI:** 10.1101/2020.04.15.042101

**Authors:** Nathalie Béchon, Alicia Jiménez-Fernández, Jerzy Witwinowski, Emilie Bierque, Najwa Taib, Thomas Cokelaer, Laurence Ma, Jean-Marc Ghigo, Simonetta Gribaldo, Christophe Beloin

## Abstract

The Negativicutes are a clade of Firmicutes that have retained the ancestral diderm character and possess an outer membrane. One of the best studied Negativicute, *Veillonella parvula*, is an anaerobic commensal and opportunistic pathogen inhabiting complex human microbial communities, including the gut and the dental plaque microbiota. Whereas adhesion and biofilm capacity of *V. parvula* is expected to be crucial for its maintenance and development in these environments, studies of *V. parvula* adhesion have been hindered by the lack of efficient genetic tools to perform functional analyses in this bacterium. Here, we took advantage of a recently described naturally transformable *V. parvula* isolate, SKV38, and adapted tools developed for the closely related *Clostridia spp*. to perform random transposon and targeted mutagenesis to identify *V. parvula* genes involved in biofilm formation. We show that type V secreted autotransporters -typically found in diderm bacteria-are the main determinants of *V. parvula* auto-aggregation and biofilm formation, which compete with each other for binding either to cells or to surfaces, with strong consequences on *V. parvula* biofilm formation capacity. We also show that inactivation of the gene coding for a poorly characterized metal-dependent phosphohydrolase HD domain protein conserved in the Firmicutes and their closely related diderm phyla inhibits autotransporter-mediated biofilm formation. This study paves the way for further molecular characterization of *V. parvula* interactions with other bacteria and the host within complex microbiota environments.

## INTRODUCTION

Negativicutes are atypical and poorly studied Firmicute lineages displaying an outer envelope with lipopolysaccharide (1). Among the Negativicutes, *Veillonella* spp. are anaerobic diderm cocci that commonly inhabit the human and animal microbiota. One of their best studied species, *Veillonella parvula* (2), is a natural inhabitant of multiple different microbiota, including the human gut (3, 4). *V. parvula* is considered a commensal organism, and proposed to play a role in the development of immunity through its capacity to colonize the infant gut (5, 6). It is a key early colonizer of the dental plaque during the establishment of sessile microbial communities called biofilms (7), promoting multi-species growth and playing a central role in the metabolism of community members through lactic acid consumption (8). However, *V. parvula* is also described as an opportunistic pathogen and has been associated with diverse infections, including osteomyelitis, endocarditis, spondylodiscitis, abscesses as well as systemic infections (9–13).

The importance of *V. parvula* in the development of microbial community spurred our interest in identifying the determinants of its adhesion and biofilm formation capacities. Moreover, considering the presence of an outer membrane in this atypical Firmicute, we wondered whether *V. parvula* uses known diderm or monoderm biofilm determinants, or rather currently undescribed adhesion factors. We previously studied *V. parvula* DSM2008 as a model diderm Firmicute strain (14) to investigate its outer membrane (OM) protein composition and detected 78 OM proteins, thirteen of which being potential adhesins belonging to the type V family of secreted autotransporter proteins (T5SS) (15). Autotransporter proteins are specifically found in diderms and all share common structural and functional features: a Sec-dependent signal peptide, a passenger domain providing the protein function, and an outer-membrane β-barrel domain that allows secretion of the passenger domain (16). However, the challenge of genetic manipulation in *V. parvula* DSM2008 severely limited the study of these adhesins in this strain.

Here, we have sequenced and annotated the genome of *V. parvula* SKV38, a recently isolated, naturally transformable and genetically amenable strain (17). We adapted and developed genetic tools for this organism, permitting random and site directed mutagenesis, plasmid complementation and controlled expression using an inducible promoter. This enabled us to identify and characterize factors involved in *V. parvula* biofilm formation. We found that the main *V. parvula* biofilm modulating determinants are T5SS adhesins, i.e. typical diderm determinants. We also showed that a locus encoding a metal-dependent phosphohydrolase HD domain protein is involved in biofilm formation, similarly to what was shown in the prototypical monoderm *Bacillus subtilis* (18). Therefore, our results demonstrate that diderm Firmicutes use a mixture of diderm/monoderm factors to modulate their ability to engage into biofilm lifestyle, supporting the idea that monoderm and diderm molecular systems could have co-evolved in these atypical Firmicutes.

## RESULTS

### PacBio sequencing of the genetically amenable *V. parvula* SKV38 strain

We sequenced the *V. parvula* SKV38 whole genome using PacBio sequencing and obtained 338,310 subreads with a mean length of 9,080 bp (Figure S1A). Assembly was performed with Canu and no gaps or drops of coverage was detected based on sequana_coverage output (Figure S1B) (19). One contig of 2,146,482 bp was generated after assembly closely matching the genome size (2.1422 Mbp) and GC content (38.7%, expected 38.6%) of the reference *V. parvula* DSM2008 strain (see supplementary material and methods for details). PROKKA annotation of the *V. parvula* SKV38 genome detected 1,912 predicted protein-encoding open reading frame (ORF), 12 rRNA, 49 tRNA and one tmRNA. Rapid Annotation using Subsystem Technology (RAST) analysis assigned 53% of the predicted proteins to known subsystems (Figure 1A). The two-way average nucleotide identity (ANI) between SKV38 and the reference strain DSM2008 is 95.43% (Figure 1B). A maximum likelihood tree of concatenated RpoB-RpoC-InfB sequences situated SKV38 strain among the Negativicutes, within the *V. parvula* clade (Figure 2).

**Figure 1:**
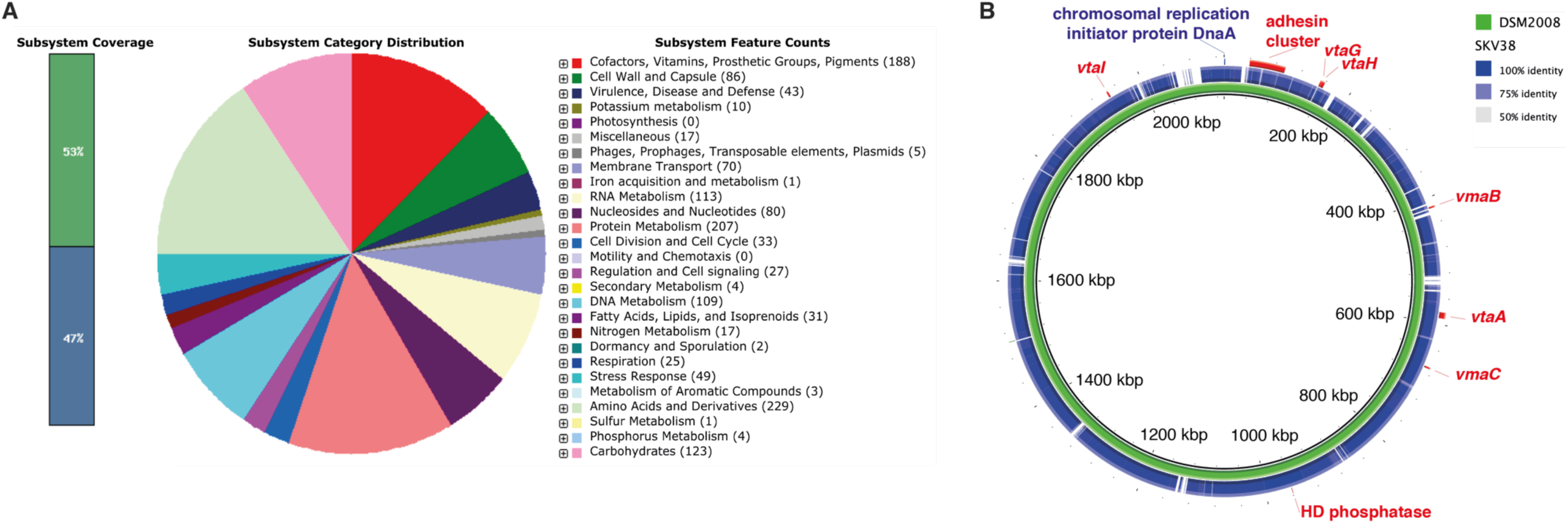
V. parvula *SKV38 genome analysis*. **A**. RAST functional annotation of the *V. parvula* SKV38 genome. Left panel: Subsystem coverage. Green color indicates the percentage of predicted proteins assigned to known subsystems, blue those that were not. Center and right panels: Subsystem category distribution: the number of proteins assigned to each category is depicted on the right panel. The analysis was performed on https://rast.theseed.org server (70–72) **B**. Comparison of the genetic identity between *V. parvula* SKV38 and the reference strain, DSM2008. The position of genes coding for potential biofilm determinants discussed in this study are indicated in the *V. parvula* genome.

**Figure 2:**
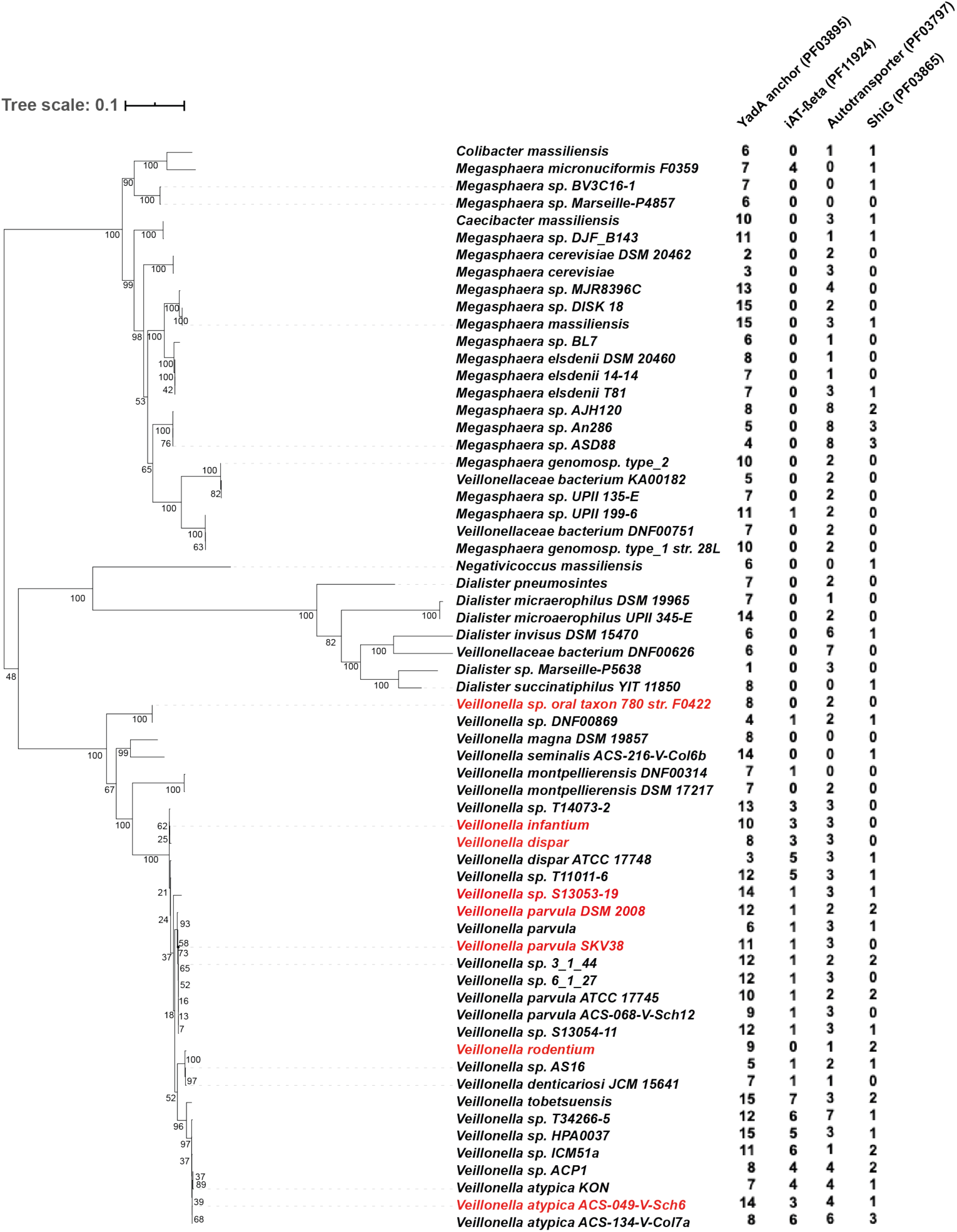
Phylogenetic tree of Veillonellaceae and prevalence of the main type V secreted adhesins. Maximum likelihood phylogenetic tree constructed from a concatenated dataset of RpoB, RpoC, InfB, comprising 3027 amino acid positions, and rooted by other Negativicutes (see material and methods). Numbers at nodes represent bootstrap values calculated based on 100 replicates of the initial character supermatrix. The number of the four main classes of potential type V secreted adhesins queried by an HMM search is indicated for every species. The scale bar indicates the average number of substitutions per site. Species whose adhesin gene clusters were analyzed on Figure 7 are colored in red.

### Random transposon mutagenesis reveals two *V. parvula* SKV38 genes involved in biofilm formation

In order to identify genes involved in biofilm formation, we performed random transposon mutagenesis in *V. parvula* SKV38 using the pRPF215 plasmid carrying an inducible transposase and a mariner-based transposon previously used to mutagenize *Clostridium difficile* (20), a close relative of the Negativicutes. We screened 940 individual transposon mutants for biofilm formation using crystal violet staining (CV) static biofilm assay in 96-well microtiter plates and identified eight independent mutants with significant reduction in biofilm formation (Figure 3A). Whole genome sequencing localized the transposons in two loci putatively implicated in biofilm formation (Figure 3B). The most affected mutants correspond to insertions in *FNLLGLLA_00516* (seven mutants), encoding a T5SS type Vc trimeric autotransporter. One transposon mutant was inserted in *FNLLGLLA_01127*, encoding a putative HD phosphatase (Figure 3B).

**Figure 3:**
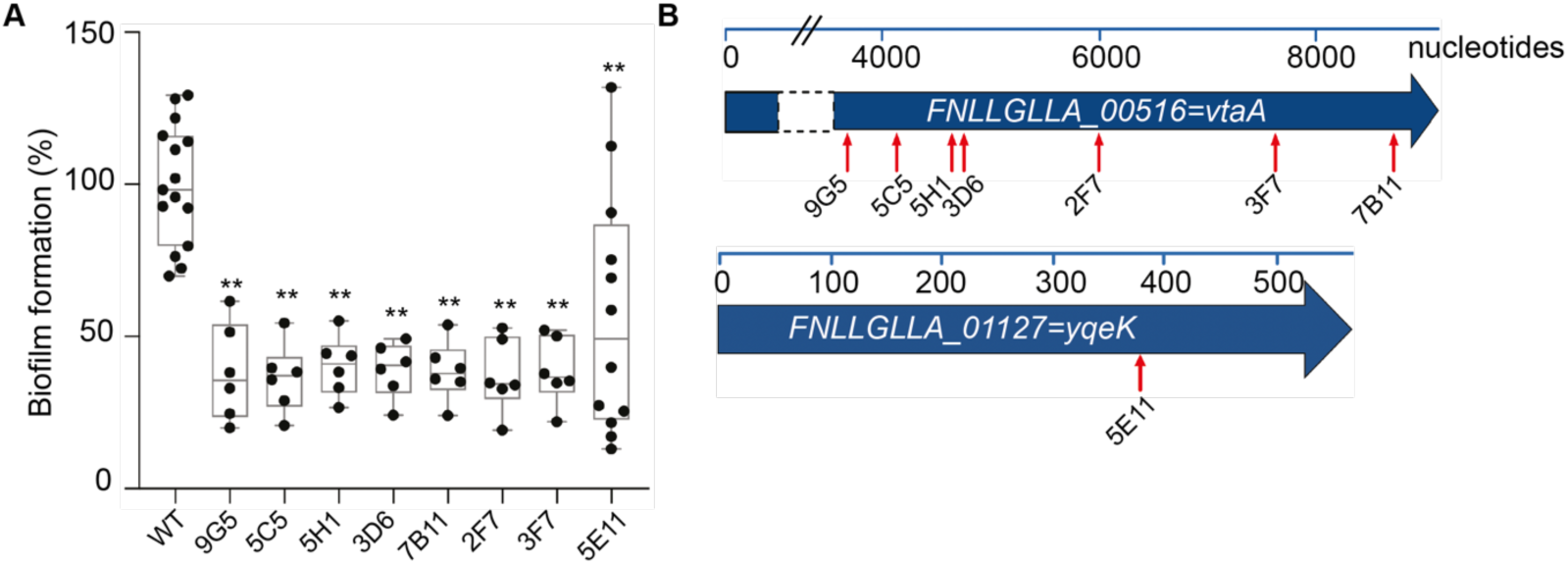
Random transposon mutagenesis in *Veillonella parvula* SKV38 led to identification of mutants with reduced biofilm formation. **A**. 96-well polystyrene plate biofilm assay after CV staining of nine transposon mutants identified by random mutagenesis grown 24h in BHILC. Mean of WT is adjusted to 100 %. Min-max boxplot of 6-15 biological replicates for each strain are represented, each replicate is the mean of two technical replicates. * p-value<0.05, ** p-value <0.005, Mann-Whitney test. **B**. Schematic representation of the transposon insertion point identified (red arrow) for the 8 transposon mutants. Blue bar represents the size of the gene in nucleotides.

### *FNLLGLLA_00516* encodes a trimeric autotransporter involved in auto-aggregation

*FNLLGLLA_00516* encodes a protein containing several domains usually identified in the T5SS type Vc trimeric autotransporters. Trimeric autotransporters are OM proteins specific of diderm bacteria that have been widely studied for their ability to bind to different surfaces or other bacteria (21). *FNLLGLLA_00516* is a homolog of *V. parvula* DSM2008 *vpar_0464*, which was shown to encode a protein detected in the OM (15). *FNLLGLLA_00516* was annotated by PROKKA as BtaF, a trimeric autotransporter identified in *Brucella suis* involved in adhesion to extracellular matrix and abiotic surfaces (22). Here, we renamed it *<u>V</u>eillonella* trimeric <u>a</u>utotransporter A (VtaA), as the first trimeric autotransporter involved in biofilm formation identified in *V. parvula* SKV38. We deleted the *vtaA* coding sequence and showed that Δ*vtaA* had no growth defect (Figure S2A) but displayed a marked reduction of biofilm formation in microtiter plate (Figure 4A). Moreover, while *V. parvula* SKV38 cultures strongly aggregated, Δ*vtaA* did not (Figure 4B and S3). We constructed the strain *pTet-vtaA*, where the chromosomal *vtaA* gene is placed under the control of a functional tetracycline/anhydrotetracycline (aTc) inducible promoter (Figure S4) and showed that its aggregation capacity and biofilm formation directly correlated with aTc concentration (Figure 4C-D), demonstrating that VtaA-mediated cell-to-cell interactions are critical for biofilm formation. Whereas the microtiter plate assay corresponds to a static biofilm assay, we also used continuous flow biofilm microfermentors to investigate the contribution of VtaA to biofilm formation in dynamic conditions. Surprisingly, Δ*vtaA* formed almost six times more biofilm than the WT strain in these conditions (Figure 4E). This suggests that auto-aggregation differentially contributes to biofilm formation in dynamic or static conditions.

**Figure 4:**
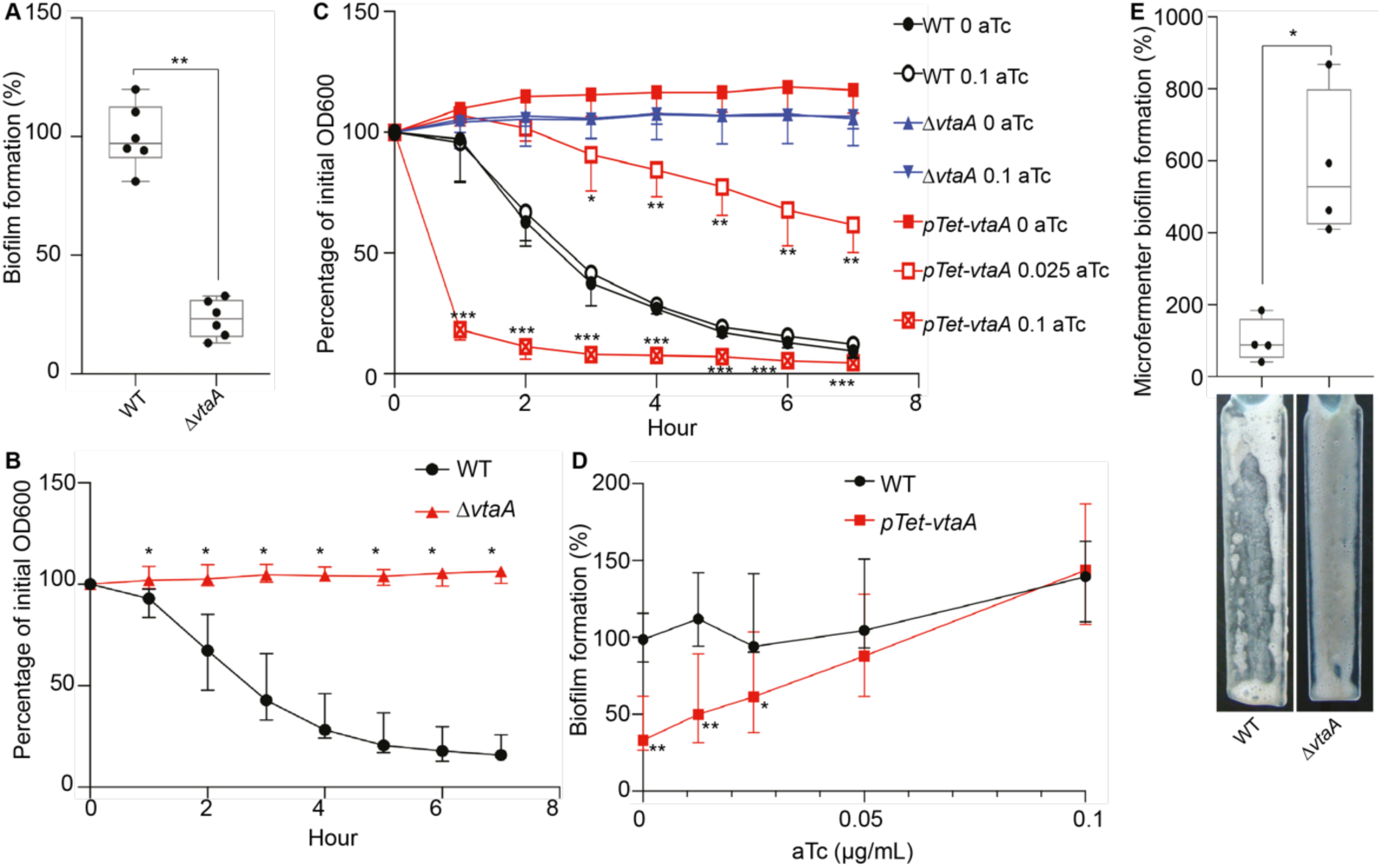
VtaA is an adhesin involved in auto-aggregation and biofilm formation. **A**. 96-well plate biofilm assay after 24h growth in BHILC. Mean of WT is adjusted to 100 %. Min-max boxplot of 6 biological replicates for each strain. * p-value<0.05, ** p-value <0.005, Mann-Whitney test between strains. **B**. and **C**. Aggregation curve in spectrophotometry cuvette of WT and Δ*vtaA* (**B**) and of an inducible *vtaA* with 0, 0.025 or 0.1 µg/mL of the inducer aTc (**C**). 100 % represent lack of aggregation, 0 % complete sedimentation of the culture. Median of 6 biological replicates, error bars represent 95% confidence interval. At each time points we computed the Mann-Whitney test between conditions. We applied Bonferroni correction for multiple testing: p-value are only considered significant if *p-value<0.004, **p-value<0.0004, ***p-value<0.00004. Indicated p-values were calculated by comparing in **B**, WT and Δ*vtaA*, and in **C**, *pTet-vtaA* without aTc and *pTet-vtaA* with different aTc concentrations. **D**. 96-well plate biofilm assay after 24h growth of an inducible *vtaA* in BHILC with different concentrations of aTc. WT without aTc is adjusted to 100 %. Median of 6 biological replicates, each replicate corresponds to the mean of two technical replicates, error bars represent 95% confidence interval. * p-value<0.05, ** p-value <0.005, Mann-Whitney test. **E**. Biofilm formation in continuous flow microfermentor on glass spatula during 48h in BHILC. WT was adjusted to 100 %. Min-max boxplot of 4 biological replicates for each strain. A picture of the spatula before resuspension is shown below each histogram bar. * p-value<0.05, Mann-Whitney test.

### *V. parvula* SKV38 encodes sixteen putative autotransporters in addition to VtaA

The strong biofilm phenotype displayed by the Δ*vtaA* mutant in microfermentor led us to suspect that additional adhesins could modulate *V. parvula* biofilm formation capacity. Indeed, searching the *V. parvula* SKV38 genome revealed multiple genes encoding autotransporters (Table 1): three Va classical monomeric autotransporters with a characteristic PFAM_PF03797 autotransporter ß-domain (renamed *<u>V</u>eillonella* <u>m</u>onomeric <u>a</u>utotransporter A to C : VmaA to C), and eight other putative Vc trimeric autotransporters with a characteristic PFAM_PF03895 YadA _anchor_domain (renamed *<u>V</u>eillonella* trimeric <u>a</u>utotransporter B to I: VtaB to I). We also identified several partial autotransporters: *FNLLGLLA_00035*, that only contains a PFAM_PF11924 Ve inverse autotransporter ß-domain but no putative α-domain that normally carries the function of the protein, and *FNLLGLLA_00036-37* and *FNLLGLLA_00040-41*, which are homologs of *V. parvula* DSM2008 *Vpar_0041* and *Vpar_0048*, respectively, and that appear to be split in SKV38 (Table 1). Among those, six autotransporters plus *FNLLGLLA_00035, FNLLGLLA_00036-37* and *FNLLGLLA_00040-41* form a potential genomic cluster coding for adhesins (Figure 1B and 5A), whereas the six others are located in different areas of the genome (Figure 1B and Figure 5B).

**Table 1:** *V. parvula* SKV38 autotransporters.

| Locus tag | PROKKA<br>Gene name | Genome position |  | Gene<br>size (kb) | Strand | Description | DSM2008<br>homolog | Name | Class |
| --- | --- | --- | --- | --- | --- | --- | --- | --- | --- |
| FNLLGLLA_00032 | prn 1 | 39,354 | 41,723 | 2,37 | forward | Autotransporter | fusion Vpar_0036-0037 | VmaA | Va |
| FNLLGLLA_00034 | btaE 1 | 42,345 | 43,754 | 1,41 | reverse | Trimeric Autotransporter: YadA like | Vpar_0039 | VtaB | Vc |
| FNLLGLLA_00035 | hypothetical<br>protein | 44,146 | 45,189 | 1,04 | forward | Autotransporter (partial) | Vpar_0040 |  | Ve |
| FNLLGLLA_00036 | hypothetical<br>protein | 45,453 | 46,883 | 1,431 | forward | none | split Vpar_0041 |  | ? |
| FNLLGLLA_00037 | omp-alpha | 46,91 | 47,878 | 969 | forward | Trimeric Autotransporter/ S-layer<br>homology domain | split Vpar_0041 |  | Vc? |
| FNLLGLLA_00038 | upaG 1 | 48,397 | 56,829 | 8,433 | forward | Trimeric Autotransporter: YadA like | Vpar_0042 | VtaC | Vc |
| FNLLGLLA_00040 | btaE 2 | 57,966 | 59,84 | 1,875 | forward | Trimeric Autotransporter: YadA like<br>(Partial) | split Vpar_0048 |  | ? |
| FNLLGLLA_00041 | ata 1 | 59,837 | 63,463 | 3,627 | forward | Trimeric Autotransporter: YadA like | split Vpar_0048 |  | Vc? |
| FNLLGLLA_00044 | ehaG 1 | 65,3 | 71,515 | 6,216 | forward | Trimeric Autotransporter: YadA like | Vpar_0051 | VtaD | Vc |
| FNLLGLLA_00045 | upaG 2 | 71,995 | 81,42 | 9,426 | forward | Trimeric Autotransporter: YadA like | Vpar_0052 | VtaE | Vc |
| FNLLGLLA_00046 | ata 2 | 81,941 | 91,519 | 9,579 | forward | Trimeric Autotransporter: YadA like | Vpar_0053 | VtaF | Vc |
| FNLLGLLA_00098 | btaE 3 | 151,792 | 153,522 | 1,731 | forward | Trimeric Autotransporter/ S-layer<br>homology domain | Vpar_0100 | VtaG | Vc |
| FNLLGLLA_00099 | ata 3 | 154,024 | 158,982 | 4,959 | forward | Trimeric Autotransporter/ S-layer<br>homology domain | absent | VtaH | Vc |
| FNLLGLLA_00335 | prn 2 | 414,666 | 416,888 | 2,223 | forward | Autotransporter | Vpar_0330 | VmaB | Va |
| FNLLGLLA_00516 | btaF | 581,236 | 590,358 | 9,123 | forward | Trimeric Autotransporter: YadA like | Vpar_0464 | VtaA | Vc |
| FNLLGLLA_00581 | brkA | 668,34 | 670,583 | 2,244 | forward | Autotransporter | Vpar_1322 | VmaC | Va |
| FNLLGLLA_01790 | ehaG 2 | 1,943,661 | 1,946,159 | 2,499 | reverse | Trimeric Autotransporter/ S-layer<br>homology domain | Vpar_1664 | VtaI | Vc |

**Figure 5:**
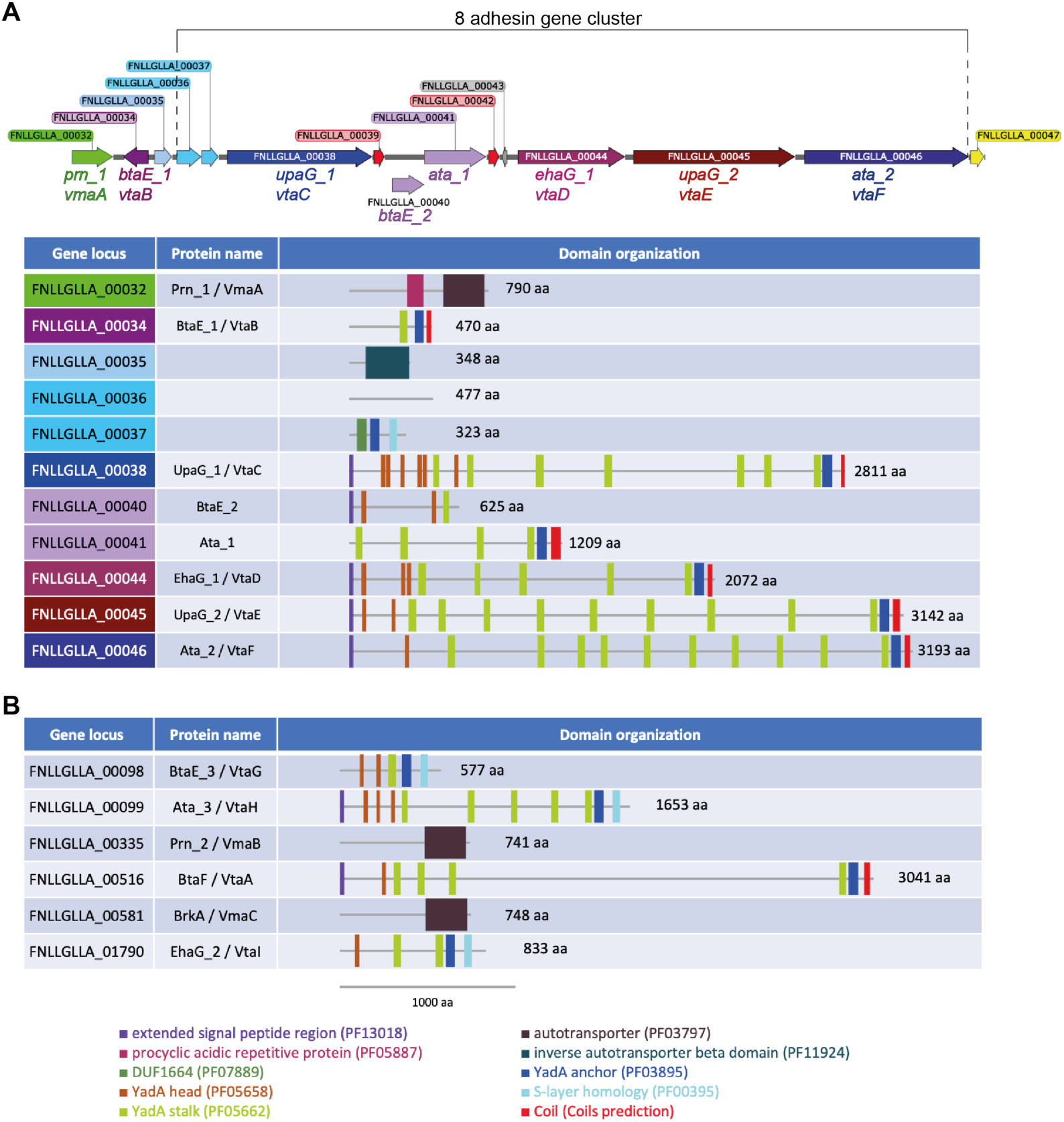
*Veillonella parvula* autotransporters domain organization. **A**. Genetic organization of the *V. parvula* SKV38 autotransporter adhesin gene cluster and the corresponding adhesin domain organization. **B**. Domain organization of the six remaining *V. parvula* SKV38 autotransporter adhesins encoded by genes located outside of the cluster. Domains were detected with the HMMER package (59), only the domains with e-values lower than 10^−3^ are shown. Presence of C-terminal coils structure was determined using the COILS program (https://embnet.vital-it.ch/software/COILS_form.html).

### The cluster of trimeric autotransporters is involved in surface binding and not aggregation

To assess the function of the potential adhesins identified in the *V. parvula* SKV38 genome, we constructed -within the cluster of adhesin genes-independent deletion mutants for the two first autotransporters (*vmaA* and *vtaB*) and a large deletion for the eight adjacent genes encoding trimeric autotransporters or partial trimeric autotransporters, hereafter called Δ*8 (*Δ*FNLLGLLA_00036* to *vtaF*). We also generated independent individual mutants for each of the six additional autotransporters located outside of the cluster. These mutants were all tested for biofilm formation and aggregation capacities. While the previously mentioned Δ*vtaA* strain was the only mutant involved in cell-to-cell interactions (Figure 6A), both Δ*vtaA* and Δ*8* led to lower biofilm formation in microtiter plate (Figure 6B and C), suggesting that the adhesins of this cluster could be involved in biofilm formation independently of cell-to-cell interactions. However, we observed no significant difference with the WT when evaluating biofilm capacity of the Δ*8* mutant in microfermentor (Figure 6D). To determine whether this was due to flow or the nature of adhesion surfaces (plastic in microtiter plate *vs*. glass in microfermentor), we used plastic microscopy coverslip slides to grow biofilms in microfermentor. Scanning electronic microscopy (SEM) showed that the WT formed a biofilm displaying filaments and protein deposits that could be part of *V. parvula* extracellular matrix, whereas Δ*vtaA* formed a much thicker biofilm, although without filaments and proteins (Figure 6E). Δ*8*, however, only poorly covered the plastic coverslip with sporadic aggregates of cells producing extracellular matrix, consistent with the reduced biofilm formation observed in microtiter plate (Figure 6E). Initial adhesion assay to glass spatula showed that both *vtaA* and Δ*8* displayed a lower percentage of initial adhesion than WT, suggesting that VtaA-mediated auto-aggregation contributed to initial adhesion while the adhesin cluster is directly involved in surface binding (Figure 6F). These two contributions were additive since a Δ*vtaA*Δ*8* double mutant showed a reduced initial adhesion on microfermentor spatula compared to either WT, Δ*vtaA* or Δ*8* (Figure 6F). In addition, Δ*vtaA*Δ*8* formed 17 times less biofilm than Δ*vtaA* in microfermentor, indicating that in the absence of VtaA, the adhesins encoded by some of these eight genes strongly promote mature biofilm formation in microfermentor (Figure 6D).

**Figure 6:**
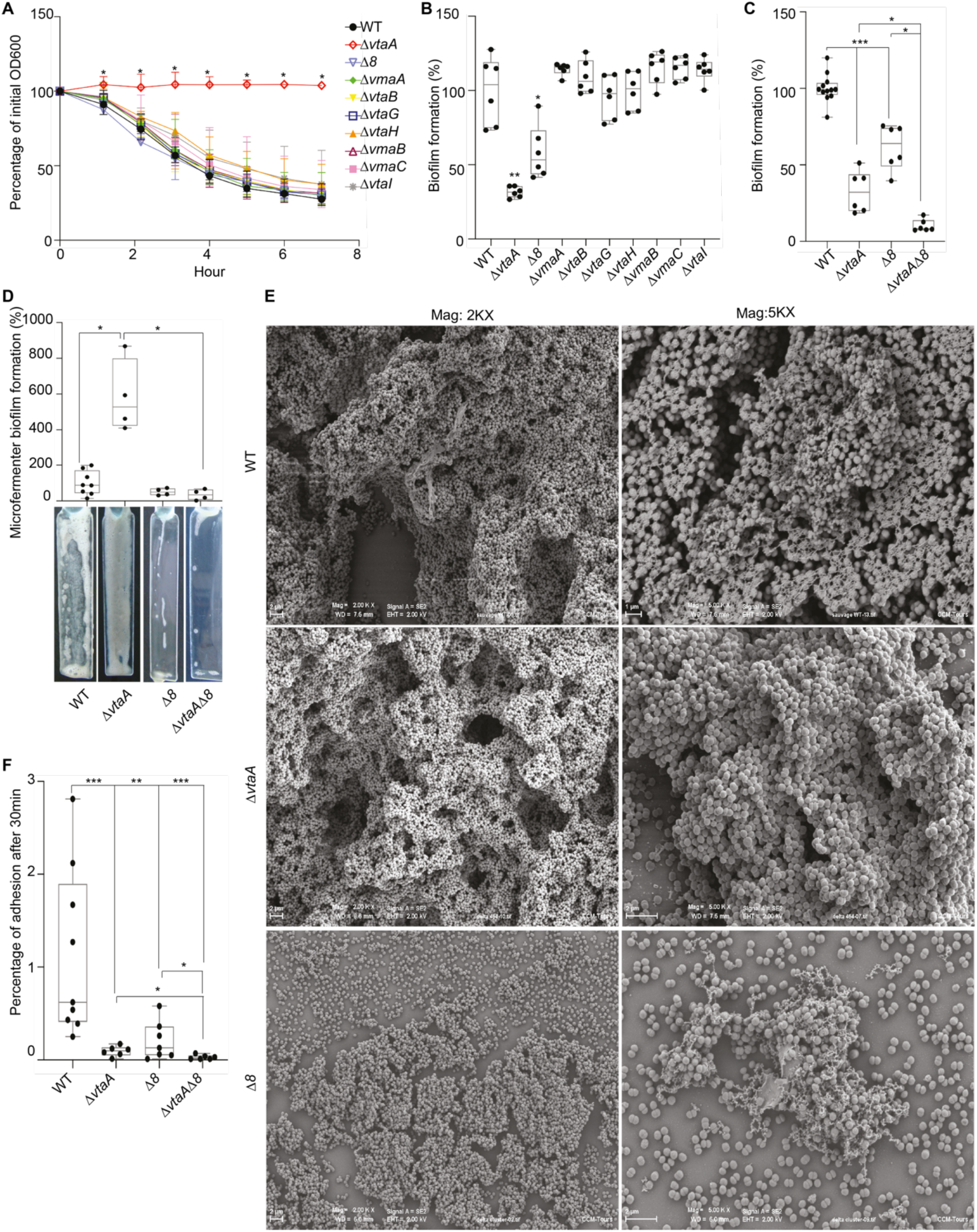
A cluster of eight trimeric autotransporters is involved in surface binding. **A**. Aggregation curve in spectrophotometry cuvette. 100 % represent lack of aggregation, 0 % complete sedimentation of the culture. Median of 6 biological replicates, error bars represent 95% confidence interval. * Mann-Whitney test, corrected for multiple testing with Bonferroni correction: significance is achieved if p-value < 0.007. **B**. and **C**. 96-well plate biofilm assay after 24 h growth in BHILC. Mean of WT is adjusted to 100 %. Min-max boxplot of 6 biological replicates for each strain, each replicate is the mean of two technical replicates. In **B**. we applied a Mann-Whitney; * p-value<0.05, ** p-value <0.005. In **C**. we applied Bonferroni correction for multiple testing: tests were called significant only if p-value<0.01: * p-value<0.01, ** p-value <0.001, *** p-value <0.0001. **D**. Biofilm formation in continuous flow microfermentor on glass spatula during 48h in BHILC. WT was adjusted to 100 %. Min-max boxplot of 4 biological replicates for each strain. * p-value<0.05, Mann-Whitney test. A picture of spatula before resuspension is shown for each mutant on the right. **E**. Scanning electronic microscopy (SEM) of biofilms grown under continuous flow of BHILC in microfermentor on plastic microscopy slide, magnified 2K or 5K times. **F**. Initial adhesion on glass spatula. Percentage of CFU that adhered to the spatula controlled by the number of CFU of the inoculation solution. Min-max boxplot of 6-9 replicates per strain is represented. * p-value<0.05, ** p-value <0.005, *** p-value <0.0005, Mann-Whitney test.

Taken together, these results demonstrate the differential contribution of VtaA and part of the cluster of adhesin to *V. parvula* SKV38 adhesion and highlight the existence of potential interference mechanisms between them.

### Prevalence of large adhesin clusters in *Veillonellaceae* genomes

The involvement of T5SS adhesins in the modulation of *V. parvula* biofilm formation prompted us to broaden our analysis and determine the prevalence of these adhesins in the *Veillonellaceae*. We plotted the results of a hidden Markov Model (HMM) search for four major types of bacterial domains found in T5SS adhesins, namely autotransporter (PF03797) typical of type Va transport, ShlB (PF03865) typical of two-partner system Vb, YadA_anchor (PF03895) typical of trimeric autotransporter type Vc transport, and iAT-ß typical of inverted autotransporter Ve (PF11924) (Figure 2). Trimeric autotransporters Vc are the most abundant potential adhesins in *Veillonellaceae*, followed by Va classical autotransporters, while type Vb two-partner system and Ve inverted autotransporters are only sporadic. Despite this trend, the numbers of each of these classes varied strongly within the same genus or even the same species of *Veillonellaceae*, indicating an important plasticity of the autotransporters repertoire.

Large clustering of potential adhesin encoding genes observed in SKV38 strain and defined as a group of two or more adhesin-coding genes immediately upstream of a conserved rRNA locus, is to our knowledge a peculiar genomic character. We found no evidence of the existence of this specific adhesin locus outside the *Veillonella* genus. We selected eight *Veillonella* strains to study more precisely the evolution of the adhesin cluster, including SKV38 and DSM2008. The trimeric autotransporter adhesins seem to evolve dynamically with numerous domain swaps, duplications and reductions of gene copies, likely through homologous recombination (Figure 7). Duplications and deletions could be eased by the presence of short ORFs annotated as hypothetical proteins presenting a high degree of sequence identity. The most basal strain in the *Veillonella* phylogeny has a minimal cluster of only three adhesins genes. Throughout the *Veillonella* genus, the size of the cluster is very variable with a minimal form in *V. atypica*, with only two adhesins.

**Figure 7:**
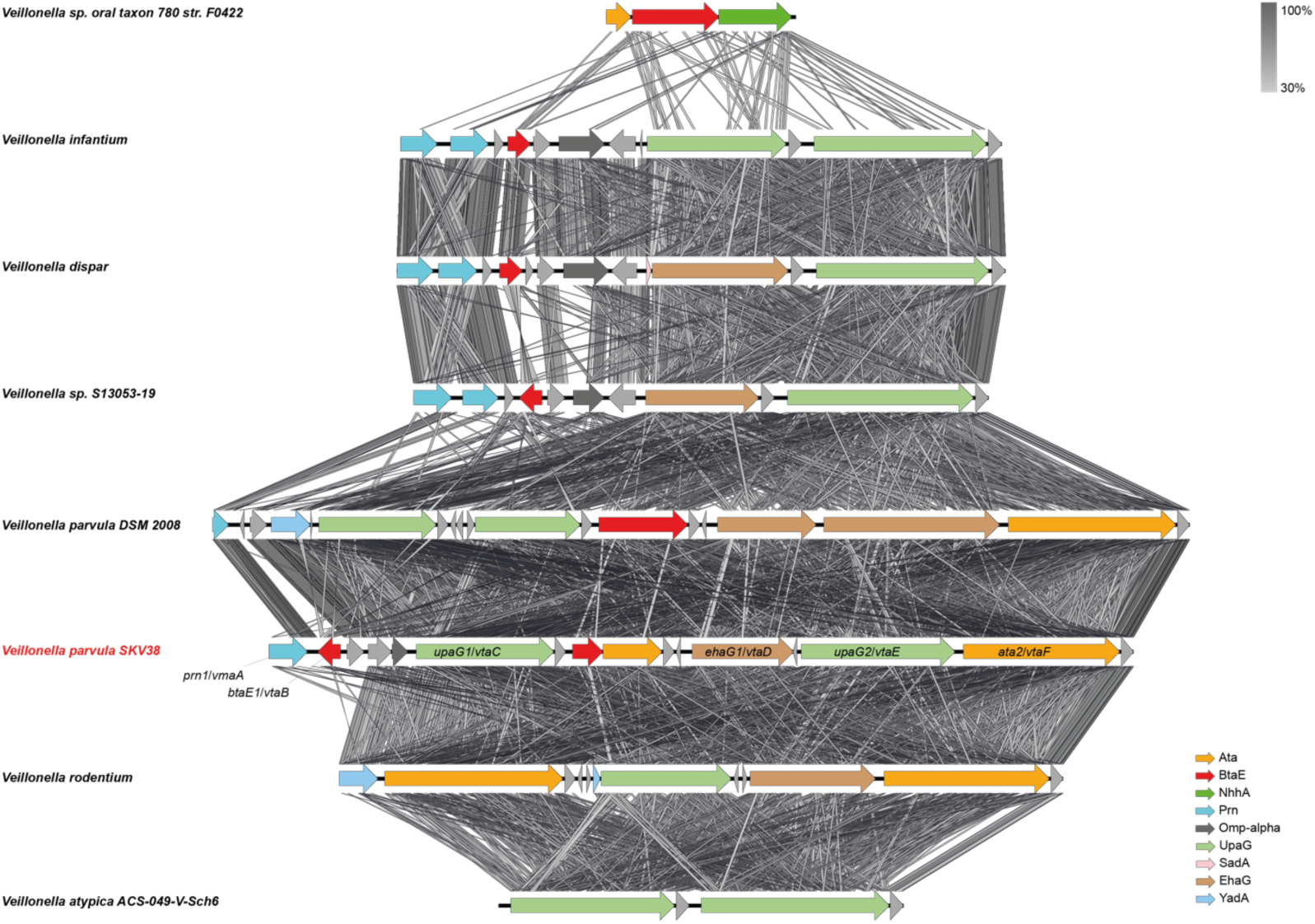
Synteny of the adhesin gene cluster in a selection of *Veillonella* species. The synteny of the proteins of the cluster between the closest relatives was assessed using EasyFig (64). Oblique lines between genes represent tblastx identities (program parameters: maximum e-value of 10^12^, minimum length of 30, minimum identity of 30). The *V. parvula* SKV38 strain used in this study is presented in red. The functional genes of the cluster are indicated.

### FNLLGLLA_01127 encodes an HD phosphatase that inhibits biofilm formation

In addition to genes encoding potential T5SS proteins, we also identified a transposon mutant in *FNLLGLLA_01127*, encoding a protein of the HD phosphatase superfamily (Figure 2). The *FNLLGLLA_01127* gene is homologous to YqeK, a putative phosphatase required for pellicle formation and the development of biofilm in *B. subtilis* (18). A *FNLLGLLA_01127* deletion mutant (Δ*1127*) showed a moderate growth defect (Figure S3AB) and a moderate 1.5-fold decrease in biofilm formation in microtiter plate after correcting for the growth defect (Figure 8A). This mutant also displayed a slightly faster aggregation rate than the WT during early time points (Figure 8B). The strongest phenotype of this mutant was detected in microfermentor with a 9-fold increase in biofilm formation compared to WT (Figure 8C), which was reduced by expressing *FNLLGLLA_01127* gene *in trans* (plasmid *p1127*) (Figure 8D). Scanning electronic microscopy showed that Δ*1127*, similarly to Δ*vtaA*, formed a thick layered biofilm, although with fewer filaments and protein deposits compared to WT (Figure 8E). However, contrary to Δ*vtaA* or Δ*8* mutants, Δ*1127* showed no defect in initial adhesion to a glass spatula (Figure 8F). Interestingly, a Δ*1127*Δ*8* double mutant formed almost 20 times less biofilm than Δ*1127* in microfermentor (Figure 8C), suggesting that at least some of the autotransporters of the cluster were necessary for Δ*1127* observed strong biofilm formation in microfermentor.

**Figure 8:**
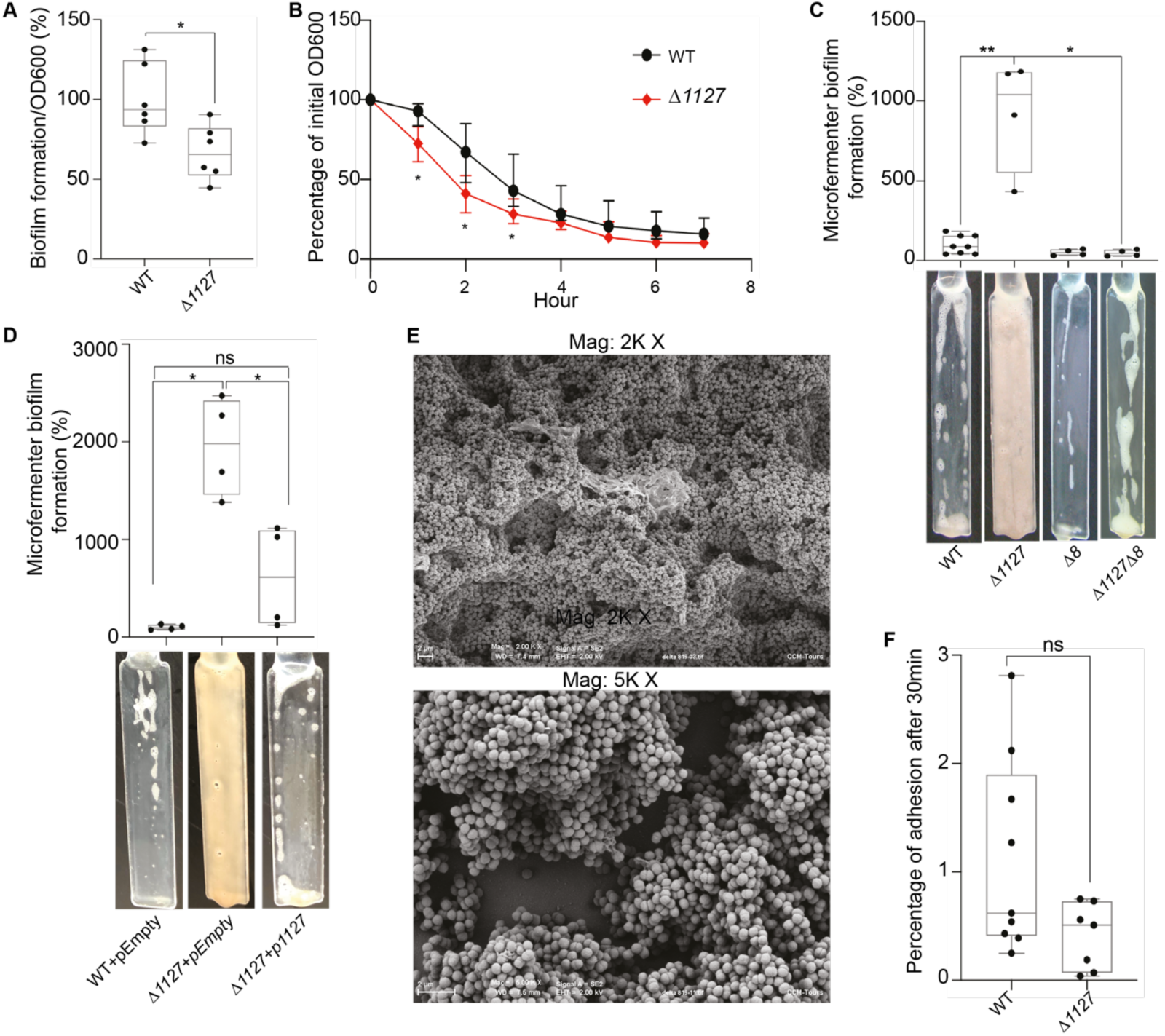
*FNLLGLLA_01127* represses biofilm formation in microfermentor. **A**. 96-well plate biofilm assay after 24 h growth in BHILC corrected by OD_600_ after 24 h growth in plate. Mean of WT is adjusted to 100 %. Min-max boxplot of 6 biological replicates for each strain, each replicate is the mean of two technical replicates. * p-value < 0.05, Mann-Whitney test. **B**. Aggregation curve in spectrophotometry cuvette. 100 % represent lack of aggregation, 0 % complete sedimentation of the culture. Median of 6 biological replicates, error bars represent 95% confidence interval. * Mann-Whitney test, corrected for multiple testing with Bonferroni correction: significance is achieved if p-value<0.007. **C**. Biofilm formation in continuous flow microfermentor on glass spatula during 48h in BHILC. Mean of WT is adjusted to 100 %. Min-max boxplot of 4 biological replicates for each strain. * p-value < 0.05, ** p-value<0.005, Mann-Whitney test. A picture of a spatula before resuspension is shown for each strain below the histogram. **D**. Biofilm formation in continuous flow microfermentor on glass spatula during 48h in BHILC+chloramphenicol. Mean of WT+pEmpty is adjusted to 100 %. Min-max boxplot of 4 biological replicates for each strain. * p-value < 0.05, Mann-Whitney test. A picture of a spatula before resuspension is shown for each strain below the histogram. **E**. Scanning electronic microscopy of Δ*1127* biofilm grown under continuous flow of BHILC in microfermentor on a plastic microscopy slide. Magnification 2K and 5K. **F**. Initial adhesion on glass spatula. Percentage of CFU that adhered to the spatula in 30 min controlled by the number of CFU of the inoculation solution. Min-max boxplot of 6-9 replicates per strain. * p-value<0.05, Mann-Whitney test.

## DISCUSSION

Originally described as a social organism mostly living in biofilm communities (8), *Veillonella* is a known bacterial member of multiple human microbiota. Although biofilm formation and adhesion are likely important in these niches, molecular studies in *Veillonella* have been hindered by the lack of efficient genetic tools. Here, we used genetics tools adapted from *Clostridia* to characterize factors promoting biofilm formation in a naturally competent *Veillonella parvula* isolate.

We identified a T5SS type Vc trimeric autotransporter, FNLLGLLA_0516 (VtaA), as an important biofilm factor promoting *V. parvula* SKV38 auto-aggregation. In addition to Hag1, a YadA-like autotransporter identified from the related species *V. atypica* involved in interspecies interactions (23), VtaA represents the first *Veillonella* protein involved in abiotic surface adhesion and auto-aggregation in diderm Firmicutes. Beyond the potential impact on *Veillonella* niche colonization, aggregation capacity contributes to bacterial protection from environmental stresses or host responses (24), promotion of host colonization (25), or pathogenesis (26). VtaA is homologous to *Brucella suis* trimeric autotransporter BtaF. However, while *B. suis* BtaF promotes biofilm formation *in vitro*, it was not shown to promote aggregation (22), suggesting that these two proteins have different functions.

In diderm bacteria such as *E. coli*, <u>s</u>elf-<u>a</u>ssociating <u>a</u>utotransporters (SAATs) from the type Va family and type Vc trimeric autotransporters were shown to contribute to biofilm formation through their self-recognition properties (27–33). However, in *V. parvula* VtaA-mediated auto-aggregation either promoted (plastic surface and static conditions) or strongly impaired (glass surface and continuous-flow conditions) biofilm formation depending on the model used. We hypothesize that under continuous flow, VtaA-mediated aggregates are more sensitive to flow than individual cells and that adhesion to surfaces or to the biofilm extracellular matrix might be more important than cell-to-cell interactions.

Interference between cell surface structures is a well-described mechanism by which bacteria modulate their adhesion properties. In *E. coli*, multiple structures, such as chaperone-usher fimbriae, LPS O-antigen or capsules, interfere with the self-recognizing autotransporter Ag43 though unknown mechanisms (34–37). Therefore, it is possible that in *V. parvula* VtaA could compete with other adhesins through steric hindrance or competition for membrane export and thus limit biofilm formation under continuous-flow conditions. Consistently, Δ*vtaA* enhanced biofilm formation in microfermentor was dependent on the presence of eight genes of the cluster of trimeric autotransporters, suggesting a competition between VtaA and adhesin(s) of this cluster. Moreover, we noticed that both VtaA and the 8-gene cluster are necessary for full initial adhesion to glass spatula in an independent manner, suggesting that any competition between them only arises later on, during continuous-flow cultures. The exact contribution of these different trimeric autotransporters to biofilm formation and their interplay with VtaA will require further characterization.

Analysis of *V. parvula* SKV38 genome revealed the presence of seven other potential full-length autotransporters, but no other types of classical diderm adhesins. None of them appeared to be involved in cell-to-cell interactions or biofilm formation on abiotic surfaces, and their function remains unclear. As *V. parvula* is present in different microbiota, it is expected that a large arsenal of adhesion factors might be necessary to adhere under different mechanical constraints and on different surfaces, such as tooth enamel or various epithelia. Moreover, *Veillonella* is known to co-aggregate with *Streptococci* (38–40), that produce *Veillonella* favored carbon source, lactate (41), and they were shown to specifically co-aggregate with *Streptococci* and *Actinomyces* strains from the same microbiota, showing that co-aggregation could have strong implication in niche colonization (42). *V. parvula* and other *Veillonella* are also associated to different opportunistic infections and the contribution of their adhesins to pathogenicity remains to be addressed. Finally, some autotransporters have been shown to carry non-adhesive functions, including protease activity (43), but we detected no classical protease domain in the *Veillonella* autotransporters.

Although we showed that genes encoding autotransporters are widespread in *Veillonellaceae* genomes, even closely related strains display different numbers, sizes and types of adhesins. For instance, *V. parvula* DSM2008 encodes three proteins with filamentous haemagglutinin domain (PF13332), whereas this class of proteins is completely absent in the closely related SKV38 genome. This suggests rapid evolutionary changes in the repertoire *Veillonella* adhesins, potentially eased by the presence of an adhesin cluster that facilitates recombination events between adjacent homologous adhesins and reduces the odds of inactivating genes involved in important cellular functions by duplication, deletion and recombination. Such a diversification might be selected for when the environmental pressure imposes to constantly adapt the adhesive properties, as it might be the case in the various niches colonized by *Veillonella*.

Trimeric autotransporters possess a characteristic YadA_anchor domain (PF03895) that can be found mainly in Proteobacteria, but also in Cyanobacteria, Verrumicrobia, Planctomycetes, Kiritimatiellaeota, Chlorobi, Synergistetes, Fusobacteria and in the diderm Firmicutes but only in Negativicutes (https://pfam.xfam.org/family/PF03895, Dec 2019 (44)). Interestingly, the YadA_anchor of all *Veillonella* trimeric autotransporters is not at the very end of the C-terminus, where it is usually found in Proteobacteria, but is pre-C-terminal, followed by either a coiled domain or a S-layer homology (SLH) domain (Figure 5). While the function of the coiled domain is unknown, in some bacteria, the periplasmic SLH domains bind peptidoglycan (45), suggesting that *Veillonella* trimeric autotransporters could be non-covalently attached to the peptidoglycan. These extra-domains after the YadA_anchor are also found in other Negativicutes (notably the extra SLH domain) and in some other diderm phyla phylogenetically related to the Firmicutes such as Synergistetes and Fusobacteria (DataSet 1). Interestingly, in addition to possessing trimeric autotransporters with an extra coiled C-terminus domain, the Fusobacteria *Streptobacillus moniliformis* ATCC14647 carries eight genes encoding unique trimeric autotransporters with an extra OmpA_family domain (PF00691) at their extreme C-terminus, a domain known to display affinity to peptidoglycan (46) (DataSet 1). This suggests that a subset of phylogenetically close diderm bacteria have evolved trimeric autotransporters integrating different peptidoglycan binding domains. Whether these domains have an impact on trimeric autotransporters function or exposure to the surface, or more generally on outer membrane stabilization is presently unknown.

Our screening also led to the identification of FNLLGLLA_01127, the homolog of *B. subtilis* YqeK, a putative phosphatase required for pellicle formation and the development of biofilm (18). *FNLLGLLA_01127/yqeK* is found in a cluster of genes (namely *obg, yhbY, proB, proA, nadD, yqeK, lytR*, and *rsfS*), whose synteny is very well conserved among Negativicutes. This cluster, or part of it, is also well conserved in almost all Firmicutes genomes we analyzed, both monoderm and diderm, as well as in members of other diderm phyla phylogenetically close to the Firmicutes, notably Deinococcus-Thermus (Figure S5 and S6, DataSet 2). *Staphylococcus aureus* YqeK was recently shown to be a nucleotidase hydrolyzing diadenosine-tetraphosphate (Ap4A) into ADP (47). In *Pseudomonas fluorescens*, an increased level of Ap4A increases cyclic-di-GMP concentration and enhances cell-surface exposure of a large adhesin LapA, thus inducing biofilm formation (48). c-di-GMP regulates biofilm formation by modulating production of a variety cell-surface appendages or exopolysaccharides in both monoderm and diderm bacteria (49–53). Interestingly, *B. subtilis* YqeK induces the *epsA-O* operon, involved in the production of biofilm matrix-forming polysaccharides (54). Deletion of *V. parvula FNLLGLLA_01127* only led to a minor decrease in biofilm formation in 96-well plate, but a strong increase in continuous-flow biofilm formation that was dependent on the presence of the cluster of trimeric autotransporters. While we cannot exclude a direct effect on the adhesins of the cluster, more likely FNLLGLLA_01127 participates to the production/regulation of an unknown exopolysaccharide, which, contrary to *B. subtilis*, would interfere with the function or exposure of the adhesins of the cluster rather than favor community development. FNLLGLLA_01127 and its homologue YqeK are both involved in biofilm formation, but the presence or absence of an outer membrane containing adhesins changes the outcome of the HD phosphatase-mediated regulation of biofilm.

In this study we have shown that classical diderm trimeric autotransporters and a potential nucleotidase, conserved both in monoderms and diderms, with possible exopolysaccharides regulatory functions are crucial for adhesion both between cells and/or to surfaces in the diderm Firmicute *V. parvula*. Our work also underscores the rapid evolution of a diverse arsenal of trimeric autotransporters in the *Veillonella* genus, both in numbers and size, probably by efficient recombination favored by gene clustering, allowing rapid adaptation to changing environments. Taken together, our results suggest a complex interplay at the surface of *V. parvula* between different cell surface structures that may have co-evolved for a long time in these atypical Firmicutes. Much remains to be discovered on the regulatory circuits controlling these adhesion factors and their role in diderm Firmicutes biology.

## MATERIALS AND METHODS

### Bioinformatic analyses

The *V. parvula* SKV38 genome was annotated using PROKKA (55) and complementary analyses were performed with RAST (56). Genetic comparison figures were generated with BRIG (57) or with SnapGene (GSL Biotech, www.snapgene.com) for visualization of genetic organization. The SKV38 annotated genome sequence was deposited in the ENA (European Nucleotide Archive) under the accession number ERZ1303164. Average Nucleotide Identity (ANI) was calculated using the following web tool http://enve-omics.ce.gatech.edu/ani/ (58). For protein domain visualization, PFAM domains (pfam.xfam.org, Pfam 32.0. (44)) were detected using HMMER (59). Domains with an e-value lower than 10^−3^ were kept and, in case of overlapping domains, the domain having the best e-value was kept. Presence of C-terminal coils structure was determined using the COILS program (https://embnet.vital-it.ch/software/COILS_form.html) (60).

For phylogenetic analyses, a local databank of the currently available 133 Negativicutes genomes was assembled and searched for InfB, RpoB and RpoC protein sequences using HMMER (59). Sequences were aligned using MAFFT (61), trimmed using BMGE with default parameters (62), concatenated, and used to generated a tree using IQTREE (63) with the LG+R5 model and ultrafast bootstrap approximation with 1000 replicates of the original dataset. Only the Veillonellaceae part of the tree was explored, the rest serving as an outgroup to root the tree. Different adhesin families were searched in the same local databank, using HMMER (59) and plotted onto the tree. *Veillonella* adhesin cluster synteny was analyzed using Easyfig for eight selected species (64) using tblastx.

### Strains and growth conditions

Bacterial strains and plasmids are listed in Table 2. *V. parvula* was grown in either Brain Heart infusion medium (Bacto Brain Heart infusion, Difco) supplemented with 0.1 % L-cysteine and 0.6 % sodium DL-lactate (BHILC) or SK medium (10 g L^**−**1^ tryptone (Difco), 10 g L^**−**1^ yeast extract (Difco), 0.4 g L^**−**1^ disodium phosphate, 2 g L^**−**1^ sodium chloride, and 10 mL L^**−**1^ 60 % w/v sodium DL-lactate, described in (17)) and incubated at 37°C in anaerobic conditions in anaerobic bags (GENbag anaero, Biomerieux, ref. 45534) or in a C400M Ruskinn anaerobic-microaerophilic station. *Escherichia coli* was grown in Lysogeny Broth (LB) (Corning) medium under aerobic conditions at 37°C. 20 mg L^-1^ chloramphenicol (Cm), 200 mg L^-1^ erythromycin (Ery) or 2.5 mg L^-1^ tetracycline (Tc) were added to *V. parvula* cultures, 100 mg L^-1^ carbenicillin (Cb) or 5 mg L^-1^ tetracycline (Tc) were added to *E. coli* cultures when needed. 100 μg L^-1^ anhydrotetracycline (aTc) was added to induce the pTet promoter unless stated otherwise. All chemicals were purchased from Sigma-Aldrich unless stated otherwise.

**Table 2:**
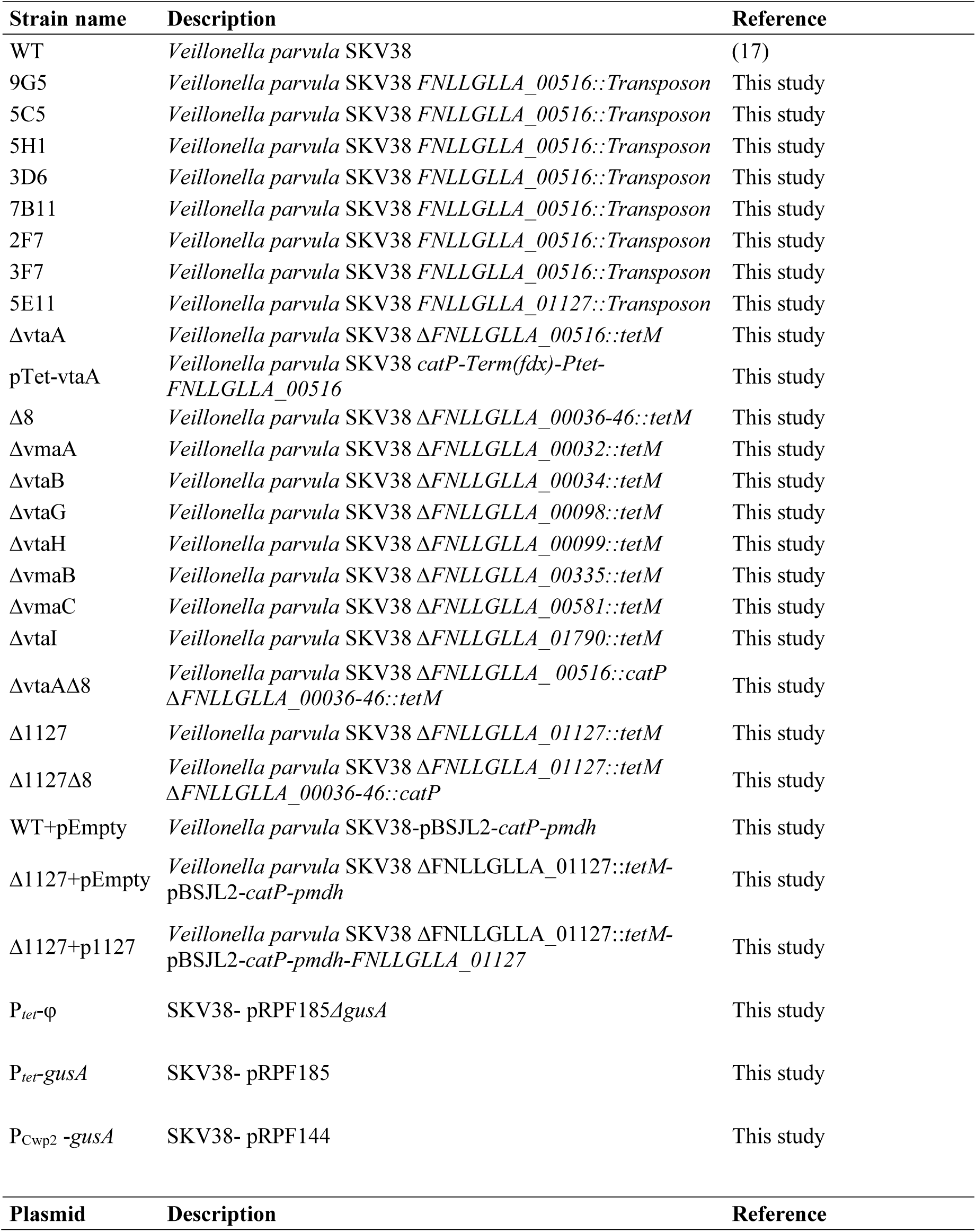
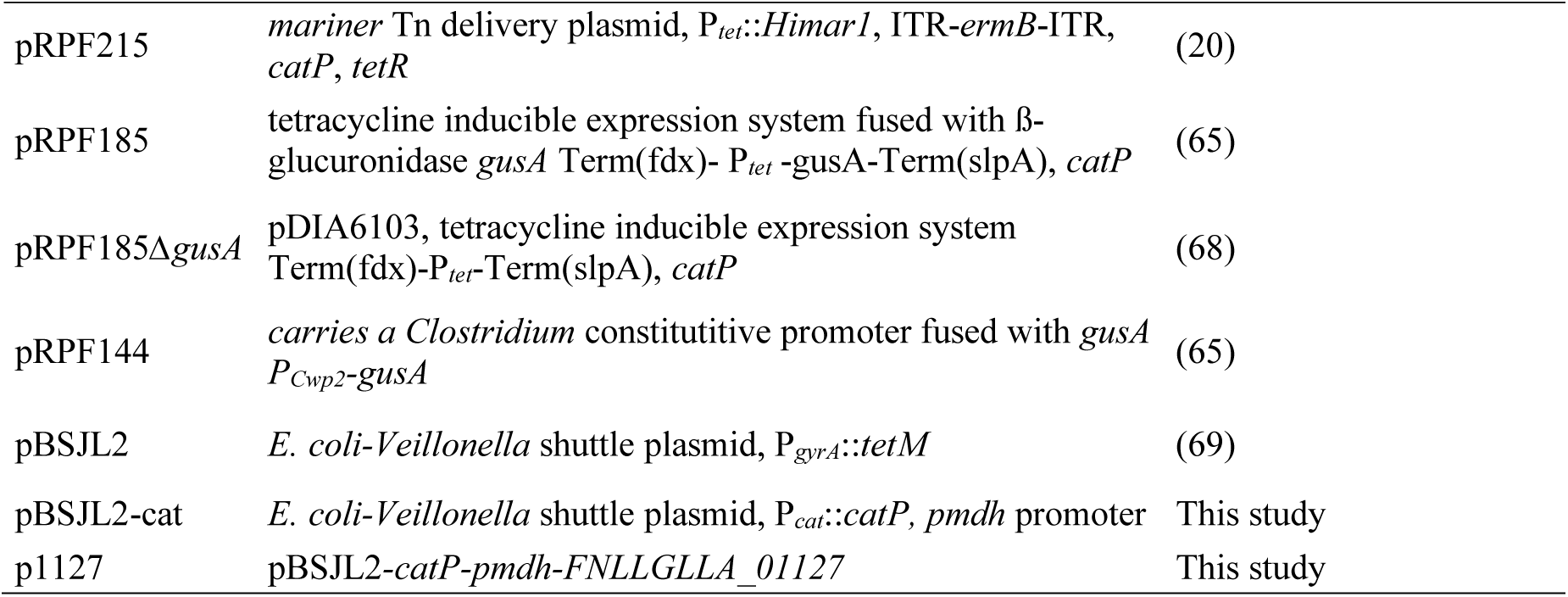
Strains and plasmids used in this study.

### Natural transformation

Cells were resuspended in 1 mL SK media adjusted to 0.4-0.8 OD_600_ and 10 µL were dotted on SK-agar petri dishes. On each drop, 0.5-1 μg plasmid or 75-200 ng μL^-1^ linear dsDNA PCR product was added, or water for the negative control. The plates were incubated 48 hours. The biomass was resuspended in 500 µL SK medium and plated on SK-agar supplemented with the corresponding antibiotic and incubated for another 48 hours. Colonies were streaked on fresh selective plates and the correct integration of the construct was confirmed by PCR and sequencing.

### Random mariner transposon mutagenesis

Plasmid pRPF215, described for *Clostridium* mutagenesis (Addgene 106377) (20) was transformed into *V. parvula* SKV38 by natural transformation and selected on Cm supplemented SK-agar medium. An overnight culture of *V. parvula* SKV38-pRPF215 in BHILC was then diluted to 0.1 OD_600_ in the same media, supplemented with aTc and grown 5 hours to induce the transposase. After induction, the culture was diluted and plated on BHILC supplemented with Ery and aTc for selection and incubated for 48 hours. From the resulting colonies, 940 were inoculated in Greiner Bio-one polystyrene flat-bottom 96-well plates (655101) and grown in BHILC supplemented with either Ery and aTc, or Cm, to confirm both the presence of the transposon and the loss of pRPF215 and then kept in 15 % glycerol at - 80°C. Selected transposon mutants were grown overnight and the genomic DNA was harvested using DNeasy blood and tissue kit (Qiagen). The genomic DNA was then sent for whole genome sequencing at the Mutualized platform for Microbiology of Institut Pasteur.

### Cloning-independent allelic exchange mutagenesis

Site-directed mutagenesis of *V. parvula* SK38 strain was performed as described by Knapp and colleagues (17). Briefly, 1-Kb regions upstream and downstream the target sequence and *V. atypica* tetracycline resistance cassette (*tetM* in pBSJL2) or catP resistance cassette from *C. difficile* (catP in pRPF185, Addgene 106367, from (65)) were PCR-amplified with overlapping primers using Phusion Flash High-Fidelity PCR Master-Mix (Thermo Scientific, F548). PCR products were used as templates in a second PCR round using only the external primers that generated a linear dsDNA with the tetracycline resistance cassette flanked by the upstream and downstream sequences. This construct was transformed into *V. parvula* by natural transformation and its integration into the genome was selected by plating on Tc or Cm supplemented medium. Positive candidates were further confirmed by a set of PCRs and sequencing around the site. Primers used in this study are listed in Table S1.

### Complementation

We replaced the tetracycline resistance gene and its *gyrA* promoter of the shuttle vector pBSJL2 by a chloramphenicol resistance gene, P_cat_:cat from pRPF185 by Gibson assembly. Briefly, the inserts and the plasmids were PCR amplified and then mixed with Gibson master mix 2x (100µL 5X ISO Buffer, 0.2 µL 10,000 U/mL T5 exonuclease (NEB #M0363S), 6.25 µL 2,000 U/mL Phusion HF polymerase (NEB #M0530S), 50 µL 40,000 U/mL Taq DNA ligase (NEB #M0208S), 87 µL dH2O for 24 reactions) and incubated at 50°C for 30-60 min.

The resulting plasmid pBSJL2-cat was digested by Fastdigest *Bam*HI (Thermo scientific) and the band obtained was purified from agarose gel using QIAquick gel extraction kit (Qiagen) to be used as linear plasmid in a second Gibson assembly. The genes and the P_mdh_ promoter of *V. parvula* SKV38 were amplified by PCR using PhusionFlash Master-mix and cloned in pBSJL2-cat using Gibson assembly. The mix was then transformed in *E. coli* DH5α and plated on LB with carbenicillin. The plasmid was harvested by miniprep using QIAprep spin miniprep kit (Qiagen) and transformed in *V. parvula* as described above.

Alternatively, the anhydrotetracycline inducible expression cassette of pRPF185, hereafter referred to as *pTet*, (Addgene 106367, (65)) was inserted along with a chloramphenicol marker right before the ATG of the target gene, following the procedure described above for cloning-independent allelic exchange mutagenesis. The functionality of *pTet* in *V. parvula* was previously verified using measurement of the aTc dependent ß-glucuronidase activity generated by the presence of pRPF185 transformed in *V. parvula* SKV38 (Figure S4).

### Biofilm formation in 96-well microtiter plates

Overnight cultures in BHILC medium were diluted to 0.05 OD_600_ and transferred to three Greiner Bio-one polystyrene flat bottom 96-well plates adding 150 μL per well. After 24 hours of static incubation, one of the three plates was resuspended by pipetting to measure OD_600_ using a TECAN Infinite-M200-Pro spectrophotometer. The two other plates were used for coloration: cultures were removed by pipetting carefully the supernatant out and biofilms fixed with 150 µL Bouin solution (HT10132, Sigma-Aldrich) for 15 minutes. Bouin solution was removed by inversion and the biofilms were washed once in water. The biofilms were stained with 150 µL crystal violet 1 % (V5265, Sigma-Aldrich) for 15 minutes without shaking, then washed in water twice and left to dry. All washes were made by flicking of the plate. After drying the plate, crystal violet was dissolved with 200 μL absolute ethanol and transferred to a clean 96-well plate for OD_620_ measurement (TECAN Infinite-M200-Pro spectrophotometer).

### Biofilm formation in microfermentor

Continuous flow non-bubbled microfermentor containing a removable spatula were used as described in (66, 67) (see https://research.pasteur.fr/en/tool/biofilm-microfermenters/). Briefly, a glass spatula was dipped in an overnight culture diluted to 0.5 OD_600_ in 15 mL BHILC for 15 minutes and returned to the fermentor. Biofilm was grown on the spatula for 48 hours at 37°C. BHILC was constantly supplied through a peristaltic pump at 4 rpm. During the last hour, the speed was increased to 10 rpm to remove planktonic bacteria. A mix of filtered 90% nitrogen/5% hydrogen/5% carbon dioxide was also constantly supplied to maintain anaerobic condition. After 48 hours of growth, the spatula was removed, and the biofilm was resuspended by vortexing in 15 mL BHILC. We measured OD_600_ of the resuspended biofilms with Smart Spec Plus spectrophotometer (BioRad).

### Aggregation curve

Overnight cultures were diluted to 0.8 OD_600_ in Brain-heart infusion (BHI) media in semi-micro spectrophotometry cuvette (Fisherbrand) and left to sediment on the bench in presence of oxygen, so no growth should occur. OD_600_ was measured every hour in a single point of the cuvette using SmartSpec spectrophotometer (BioRad).

### Initial adhesion on glass

Glass spatula from microfermentor (described above) were dipped in overnight cultures diluted to 0.5 OD_600_ in 15 mL Brain-Heart Infusion (BHI) media for 30 minutes to let bacteria adhere. The spatulas were washed once in 15 mL BHI by submersion and the adhering bacteria were resuspended in 15 mL clean BHI by vortexing. The culture used for inoculation, as well as the resuspended bacteria were serially diluted and plated on SK-agar plate for colony forming unit (CFU) counting.

### Statistical analysis

Statistical analysis was performed using either R and Rstudio software or GraphPad Prism8 (GraphPad software, Inc.). We used only non-parametric test, and when applicable corrected for multiple testing. For microfermentor experiments, 4 replicates of each condition were used. For all the other experiments, at least 6 biological replicates in at least 2 independent experiment were used. A cut-off of p-value of 5% was used for all tests. * p<0.05; ** p<0.05; *** p<0.005. For growth curve analyses, we computed the growth rate and carrying capacity of each biological replicates using Growthcurver 0.3.0 package in R and we performed a Mann-Whitney test comparing both parameters for each mutant to the corresponding WT.

## COMPETING FINANCIAL INTERESTS

The authors declare no competing financial interests.

### ACKNOWLEDGEMENTS

We thank Justin Merritt for providing *V. parvula* SKV38 strain, Bruno Dupuy and Robert P. Fagan for providing the different *Clostridium* plasmids, Pierre Simon Garcia for help with the Figure 5 preparation, Daniela Megrian Nuñez and Panagiotis Adam for the genome databank preparation and the platforms France Génomique and IBISA. We wish to acknowledge funding from the French National Research Agency (ANR) (Fir-OM ANR-16-CE12-0010), from the Institut Pasteur “Programmes Transversaux de Recherche” (PTR 39-16), from the French government’s Investissement d’Avenir Program, Laboratoire d’Excellence “Integrative Biology of Emerging Infectious Diseases” (grant n°ANR-10-LABX-62-IBEID) and from the Fondation pour la Recherche Médicale (grant DEQ20180339185). N.B. was supported by a MENESR (Ministère Français de l’Education Nationale, de l’Enseignement Supérieur et de la Recherche) fellowship. A.J.F. was supported by a PRESTIGE program from Campus France.

## AUTHORS CONTRIBUTIONS

C.B., N.B. and A.J.F. designed the experiments. N.B., A.J.F., E.B. and L.M. performed the experiments. J.W., N.T., and T.C. carried out all genomics and phylogeny analyses under the supervision of S.G. N.B., C.B. and A.J.F. wrote the paper, with contribution from S.G., J.W., T.C. and JM.G. All authors have read and approved the manuscript.

