## Supplemental M&M and Figures for "Autotransporters drive biofilm formation and auto-aggregation in the diderm Firmicute *Veillonella parvula*"

### SUPPLEMENTARY MATERIAL

#### Supplementary Material and Methods

##### Genome preparation and sequencing

*V. parvula* SKV38 genomic DNA was extracted using Qiagen genomic tip 20G kit. It was sequenced to 1,500X coverage using PacBio sequencing of one single molecule real time (SMRT) cell with no multiplexing using the V2.1 chemistry. Only one SMRT cell was used but with no multiplexing, leading to an unusually large amount of subreads: 3 Gbp, meaning about 1,500X coverage assuming a 2.1 Mbp genome. This yielded 338,310 reads with a mean subread length of 9,080 bp and N50 read length of 13,500 bp. The longest subread length is above 70 kbp. A histogram of the subread length is shown in Figure S1A. We randomly subsampled the data to avoid misassemblies keeping only 100,000 subreads, which resulted in a 430X coverage. The genome was then assembled using Canu version 1.8 (1) keeping the default parameters. In particular, subread below 1,000 bp were dropped. The error correction steps of the Canu algorithm were not tuned, keeping the parameters that control alignment seed length, read length, overlap length and error rates to their default values. We obtained one contig of about 2.167 Mbp and an additional contig of only 1,972 bp that was abandoned due to lack of supporting data and was removed by the circularization process. The resulting assembled genome was polished using Pilon (2) but no correction was required. The completeness of the candidate assembly was evaluated by running the Sequana_coverage tool (3, 4) and BUSCO version 3.0.2 (5), using the bacteria mode and the bacteria_db9 lineage-specific profile library. The completeness of the candidate assembly was assessed to be 98% using the BUSCO software (5), while the number of complete duplicated or fragmented BUSCOs remains at 0, indicative of complete assembly. Moreover, as shown in Figure S1B, no gaps or drops of coverage were detected, and alignment of all reads show that only 4% (13,028) remained unmapped and 80% of their length were below 2 kbp. The remaining reads (2000 reads) map on various species and could not be further assembled. Overall, these analyses indicate that the final genome assembly is complete and of good quality.

##### Growth curve

Overnight cultures were diluted to 0.05 $OD_{600}$ in 150µL BHILC that had previously been left in anaerobic condition overnight to remove dissolved oxygen, in Greiner flat-bottom 96-well plates. A plastic adhesive film (adhesive sealing sheet, Thermo Scientific, AB0558) was added on top of the plate inside the anaerobic station, and the plates were then incubated in a TECAN Infinite M200 Pro spectrophotometer for 24 hours at 37°C. $OD_{600}$ was measured every 30 minutes, after 900 seconds orbital shaking of 2 mm amplitude.

**Scanning electronic microscopy**

Spatula carrying Thermanox plastic coverslips (10.5 x 22 mm, Nunc Thermo Scientific) were introduced in our microfermentor system described above. After 48 hours of growth of the biofilm, the microscopy coverslips were submerged in fixation solution: volume/volume mix of glutaraldehyde 4 %/ sodium cacodylate trihydrate buffer 0.2M pH=7.4/ Ruthenium red 0.15 % and kept at 4°C. Scanning electronic microscopy was then performed by the *Laboratoire de biologie cellulaire et microscopie électronique* of the University of Tours, France.

**β-glucuronidase assay**

Overnight cultures were diluted to OD 0.05 and grown for 2-3 hours after which the inducer aTc was added. Induced cultures and controls were cultured for 3.5 hours and then ß-glucuronidase activity was assessed. 75 µl of the culture was mixed with 675 µl of Z-buffer (60 mM Na2HPO4; 40 mM NaH2PO4; 10 mM KCl; 1 mM MgSO4; 50 mM HO-CH2-CH2-SH), 50 µl of CHCl3 and 25 µl of 0.1% sodium dodecyl sulfate solution, vortexed and incubated 5 min at 37°C. Then 100 µl of 10 mM 4-nitrophenyl-β-D-glucuronide (in water) was added, and the reaction mix was incubated at 37°C until it turned visibly yellow. The reaction was stopped by adding 375 µl of 1M Na2CO3 (in water) and the absorbance at 405 nm wavelength was measured after a centrifugation. Then the β-glucuronidase activity expressed in Miller units was calculated as described previously for β-galactosidase (6).

**Light microscopy**

Cultures were grown overnight with no shaking and the aggregates at the bottom of the tubes were harvested. 1µL of these aggregates were dropped on multitest 12-well slide (MP Biomedicals glass), covered with a coverslip (24 x 60 mm coverslip, Menzel glaser) and observed using light microscopy, 1000X.

**Occurrence and synteny of FNLLGLLA_01127/YqeK**

The search for HD phosphatase (YqeK) cluster homologs was conducted as follows. A local databank containing 390 representative bacterial genomes was mined for the presence of proteins containing an HD domain (PF01966) using HMMSEARCH and the --cut_ga option. Results were then manually inspected to separate true YqeK homologues from other protein families using a combination of alignment, functional annotation, protein domains presence and phylogeny. Synteny was assessed using MacSyFinder (7) programmed to search for at least one of the following proteins: NadD (containing CTP_transf_like domain, PF01467), RsfS (containing RsfS domain, PF02410), YhbY (containing CRS1_YhbY domain, PF01985), LytR (containing LytR_cpsA_psr domain, PF03816), Obg (containing GTP1_OBG domain, PF01018), ProB (containing AA_kinase domain, PF00696) and ProA (containing Aldedh domain, PF00171), in the context of the HD containing proteins identified in the first step, with no more than eight other genes separating them. All HMM profiles were downloaded from the PFAM site (pfam.xfam.org), (Pfam 32.0 (8)) and mapped on a schematic bacterial phylogenetic tree representing only 187 cultured organisms among the 390 species of our local databank. As YqeK homologs are widespread in the Firmicutes, another local databank containing 230 representative Firmicutes genomes was queried by the MacSyFinder approach as described above. All trees were visualized with ITOL (9). Details of the results are presented in Dataset S2.

**Supplementary Table**

| Primer name | Sequence |
|---|---|
| TetM-F | agtaaaatgcaggcgagtgaag |
| TetM-R | gtggatccacaggacacaat |
| TetM-outF | agatttgaattaaagtgtaaaggagga |
| TetM-outR | tcttgttaacagcgcaattcct |
| catP-F | ggccttttgctcacatgttc |
| catP-R | cctgaagttaactatttatcaattcctgc |
| catP-chF | cgcagtatgtgacggatttc |
| catP-chR | gcttcctcgctcactgactc |
| vtaA-extF | ggctgatatgcatcccac |
| vtaA-extR | cacttacaaggcccacgact |
| vtaA-5F | gtagctttagccaatgtggc |
| vtaA-5R | cttcactcgcctgcattttactactttctcctataaactatagtaatattaatgc |
| vtaA-3F | attgtgtcctgtggatccacgagttgagaactaattttgataatagtc |
| vtaA-3R | gtatacataccgaaccgctc |
| cat-vtaA-5R | gaacatgtgagcaaaaggccactttctcctataaactatagtaatattaatgc |
| cat-vtaF-3F | gataaatagttaacttcagggagttgagaactaattttgataatagtc |
| 0036-extF | cagcctatgcaggggactac |
| 0046-extR | tctgtgccccttcatatggg |
| 0036-5F | gtcagggcggattggagatg |
| 0036-5R | cttcactcgcctgcattttacttttatcctctccattccctaatgaa |
| 0046-3F | attgtgtcctgtggatccactacgttcatactgaacattacgg |
| 0046-3R | tttgcacaatttcacggcc |
| cat-0036-5R | agaacatgtgagcaaaaggcctttatcctctccattccctaatgaa |
| cat-0046-3F | gataaatagttaacttcaggtacgttcatactgaacattacgg |
| vmaC-extF | caatccttgatcgcaacgc |
| vmaC-extR | ctacgtaatttaacgctcccc |
| vmaC-3R | tagtgttccggctaggatg |
| vmaC-3F | attgtgtcctgtggatccaccggcataatttttaattgaatattg |
| vmaC-5F | atataaaggcgctccttagg |
| vmaC-5R | cttcactcgcctgcattttactaaaaatctctctttcttacctcgc |
| vtaH-extF | ggttatggtcactaccataatgc |
| vtaH-extR | caaaatcaacatatggtcacatcc |
| vtaH-3R | cctgatctgcgatcagagc |
| vtaH-3F | attgtgtcctgtggatccaccgcggttaatagaatcgaaag |
| vtaH-5F | cgcattctaccgccctaac |
| vtaH-5R | cttcactcgcctgcattttactccggttcataacactcctcc |
| vtaB-extF | ccgttagacacattttccatgc |
| vtaB-extR | gagtcccaagtaggtcgtgc |

| | |
|---|---|
| vtaB-3R | cgtatacaattggtgcacgc |
| vtaB-3F | attgtgtcctgtggatccactaagatccctttatggtagtcctatg |
| vtaB-5F | accagacacaccgcgatac |
| vtaB-5R | cttcactcgcctgcattttactctcatcgtatacactcctttgtgtac |
| vmaA-extF | gcttctttcaattgctattaggc |
| vmaA-extR | ggtaaaagtacaagtcaacagc |
| vmaA-3R | gtacaacatgtatctcaagagg |
| vmaA-3F | attgtgtcctgtggatccacgcgaattattaatagataggagcg |
| vmaA-5F | tgcatgataataagctattccttc |
| vmaA-5R | cttcactcgcctgcattttactcataataaccctctcttattcacac |
| vtaG-extF | caatcccctgatacacgtg |
| vtaG-extR | catcatcttcagccaccgc |
| vtaG-3R | cctttgcctgcgcattac |
| vtaG-3F | attgtgtcctgtggatccacgacgtatacggttcccgtatatac |
| vtaG-5F | agctattgctgtagggtctc |
| vtaG-5R | cttcactcgcctgcattttactctctcatcttacaccctcac |
| vmaB-extF | caagtttcaaggagtactcg |
| vmaB-extR | gaataaaatattcaaaagcatctcc |
| vmaB-5F | ttcttcttgggaaatgaagttgactacagt |
| vmaB-5R | cttcactcgcctgcattttactgttatacctccttaacacttcaac |
| vmaB_3F | attgtgtcctgtggatccacttaagggtaaaaatagcagaggc |
| vmaB-3R | gaaaataatggttgagtctcag |
| vtaI-extF | cccgcatcaaagagcattgg |
| vtaI-extR | aaacatgcacgatggcccta |
| vtaI-5F | taataaactagcaggcatgcg |
| vtaI-5R | cttcactcgcctgcattttactcgtttactattcctccctacatc |
| vtaI_3F | attgtgtcctgtggatccacttacttgtcatgaattaaccttttaaac |
| vtaI-3R | caaagagcattggaatgcc |
| 1127-extF | ggtgttcgcattcacggag |
| 1127-extR | gtgcccccattatgttccg |
| 1127-5F | gtagctcaatatatggaaaacac |
| 1127-5R | cttcactcgcctgcattttactcatttattccttcccatatatatggttttc |
| 1127_3F | attgtgtcctgtggatccactggagttttatatggacgaacg |
| 1127-3R | ggtgcaactttcgacatac |
| pvtaA-3F | gcgttaacagatctgagctccttattactatagtttataggagaaagtatg |
| pvtaA-3R | ctacagaacgactcgcag |
| pvtaA-5R | aagaacatgtgagcaaaaggccattcataattctatatagcgcac |
| pvtaA-extF | ctatctattggggcagg |
| pvtaA-extR | agcattaacattatatcccgc |
| pTet-chF | cgttaacagatctgagctcc |
| pTet-chR | ccttgaattgatcatatgcgg |
| catR-pTet | tagcttgatgcagaattcgccctgaagttaactatttatcaattcctgc |

| | |
|---|---|
| cat-pTet-F | gataaatagttaacttcagggcgaattctgcatcaagc |
| pTet-R | aggagctcagatctgttaac |
| pBSJL2_ΔtetR | tgtattttatgttgttatataaatatg |
| pBSJL2_ΔtetF | ctgcaggaattcgatatc |
| cat_pBSJL2_R | catatttatataacaacataaaatacattaactatttatcaattcctgcaattcgt |
| cat_pBSJL2_F | gatatcgaattcctgcagagtgagctgataccgctcg |
| pmdh-F | caacaatcactagtggatccataccaaaattcttcaaaaaaatc |
| pmdh-R | cgttaaaacctcttttcagaaaatatg |
| mdh-1127-F | tctgaaaagaggttttaacgatgttatcgtttgatgaaatacaac |
| pBSJL2-1127-R | atattgtgtcctgtggatccttatttacaagattttaatacatcattacg |

Table S1- Primers used in this study.

#### Supplementary figures

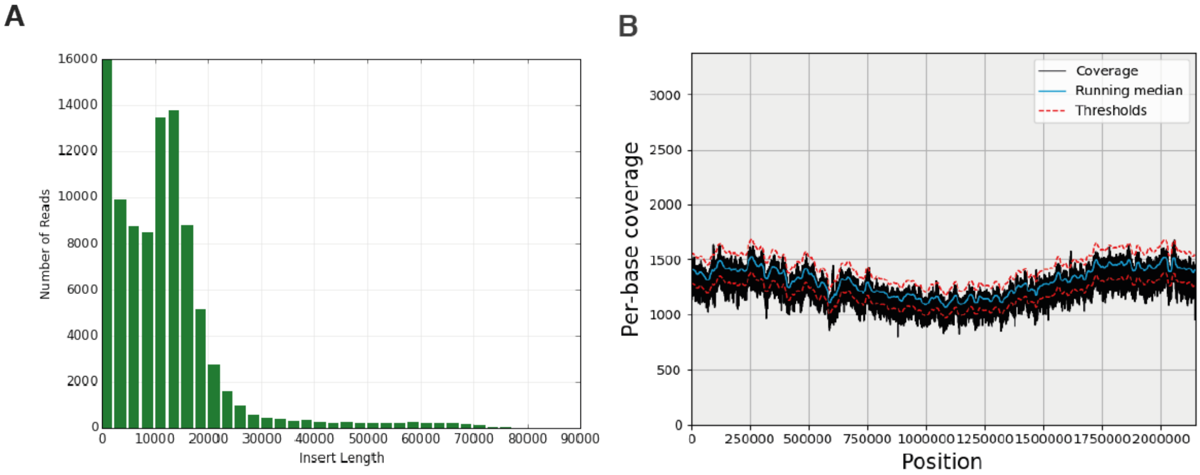


Figure S1**: PacBio Sequencing of SKV38 *V. parvula* genome. A.** Histogram of the subread length of PacBio sequencing. **B.** Coverage along the final candidate assembly of the PacBio subreads. See detailed Supplementary Material and Methods.

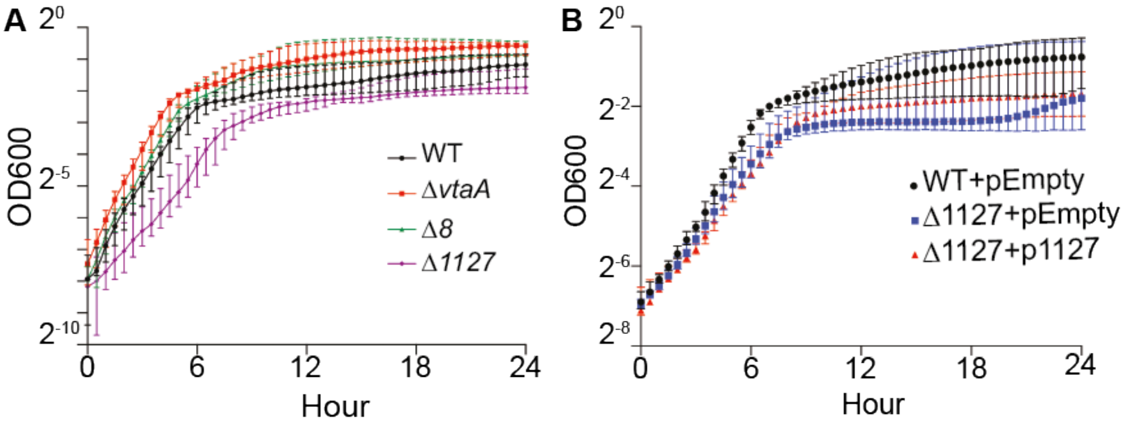


| | Growth rate | | Carrying capacity | |
|---|---|---|---|---|
| | median | p-value | median | p-value |
| WT | 0.33 | | 0.41 | |
| *ΔvtaA* | 0.39 | 0.297 | 0.65 | 0.024 |
| *Δ8* | 0.47 | 0.145 | 0.53 | 0.088 |
| *Δ1127* | 0.38 | 0.605 | 0.26 | 0.040 |

| | Growth rate | | Carrying capacity | |
|---|---|---|---|---|
| | median | p-value | median | p-value |
| WT+*pEmpty* | 0.37 | | 0.56 | |
| *Δ1127+pEmpty* | 0.57 | 0.818 | 0.21 | 0.041 |
| *Δ1127+p1127* | 0.55 | 0.589 | 0.29 | 0.026 |

Figure S2: **Growth of *V. parvula* mutants.** 24h growth curve in BHILC in 96-well plates. Each point represents the median of 6 biological replicates. Error bars represent 95% confidence interval. For each biological replicate of each strain, the growth rate and the carrying capacity were computed using R library growthcurver. The median of both parameters is indicated in the table below, as well as the p-value of a Mann-Whitney test comparing growth rate and carrying capacity of the mutant with the corresponding WT or WT+pEmpty. **A**. Selected deletion mutants. **B.** Complementation of the Δ1127 mutant. pEmpty correspond to pRPF185. p1127 corresponds to pBSJL2-catP-pmdh-FNLLGLLA_01127. See detailed Supplementary Material and Methods.

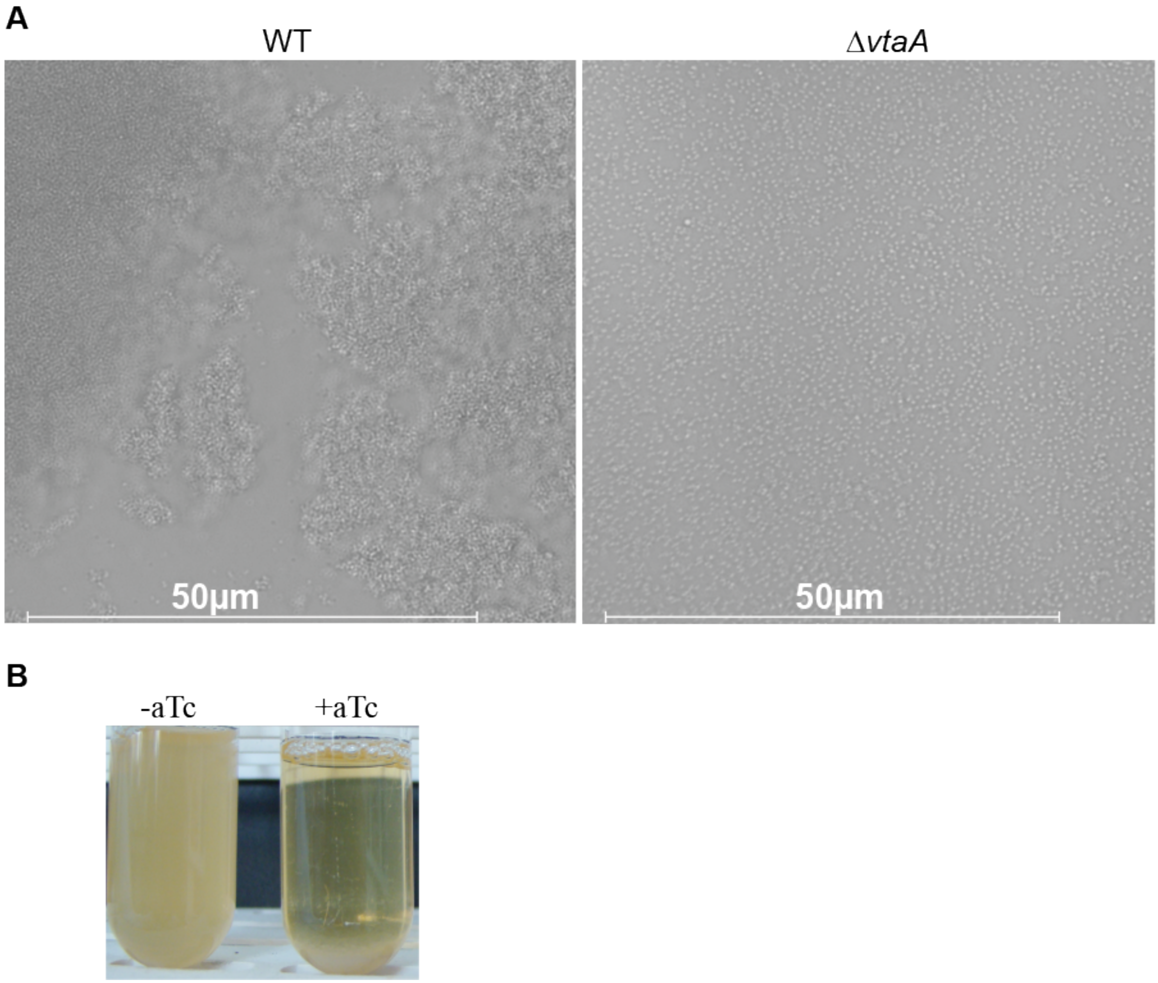


Figure S3: **VtaA mediates auto-aggregation. A.** Light microscopy of overnight cultures of WT and Δ*vtaA* grown in BHILC, 1000X. Scale bar represent 50µm. **B.** Overnight cultures of *pTet-vtaA* strain*,* where *vtaA* is under the control of the inducible pTet promoter, grown in BHILC with or without 0.1µg/mL aTc.

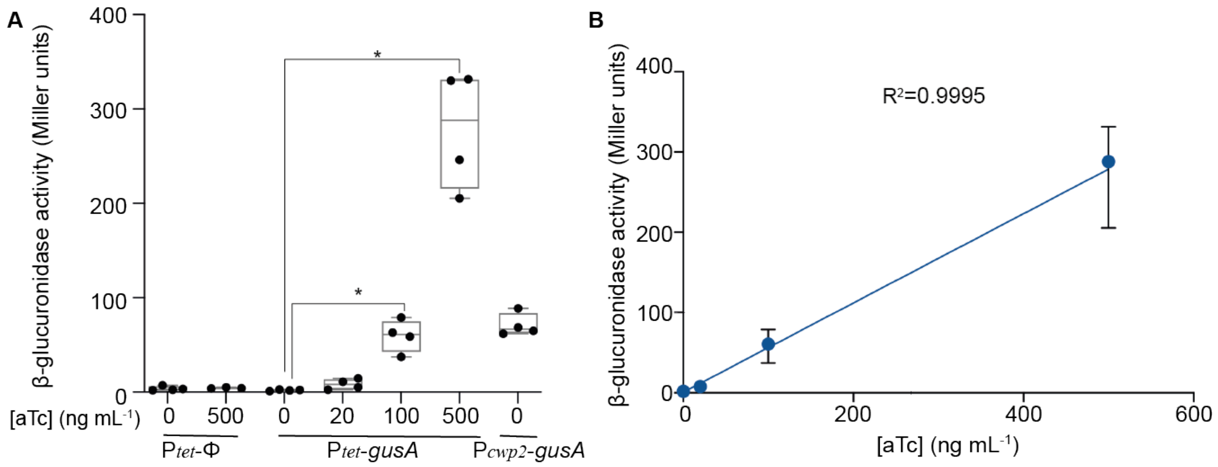


Figure S4. **Tetracycline inducible promoter (P$_{tet}$) from *Clostridium difficile* is active in *V. parvula*. A.** β-glucuronidase assay showing the expression of P$_{tet}$ fused to *gusA* under different concentrations of aTc. SKV38 was transformed with pRPF185*ΔgusA* (P$_{tet}$-φ) as a negative control, pRPF185 carrying the P$_{tet}$-*gusA* or pRPF144 carrying a copy of *gusA* under a constitutive promoter from Clostridium (P$_{Cwp2}$ -*gusA*). Min-max boxplot of 4 biological replicates. *p-value<0.05, Mann-Whitney test. **B**. ß-glucuronidase activity measured for SKV38-pRPF185 carrying P$_{tet}$-*gusA* is represented as a function of the inducer aTc concentration. Each dot is the median of 4 biological replicates, error bars represent 95% confidence interval. The coefficient of correlation $R^2$ is indicated. See detailed Supplementary Material and Methods.

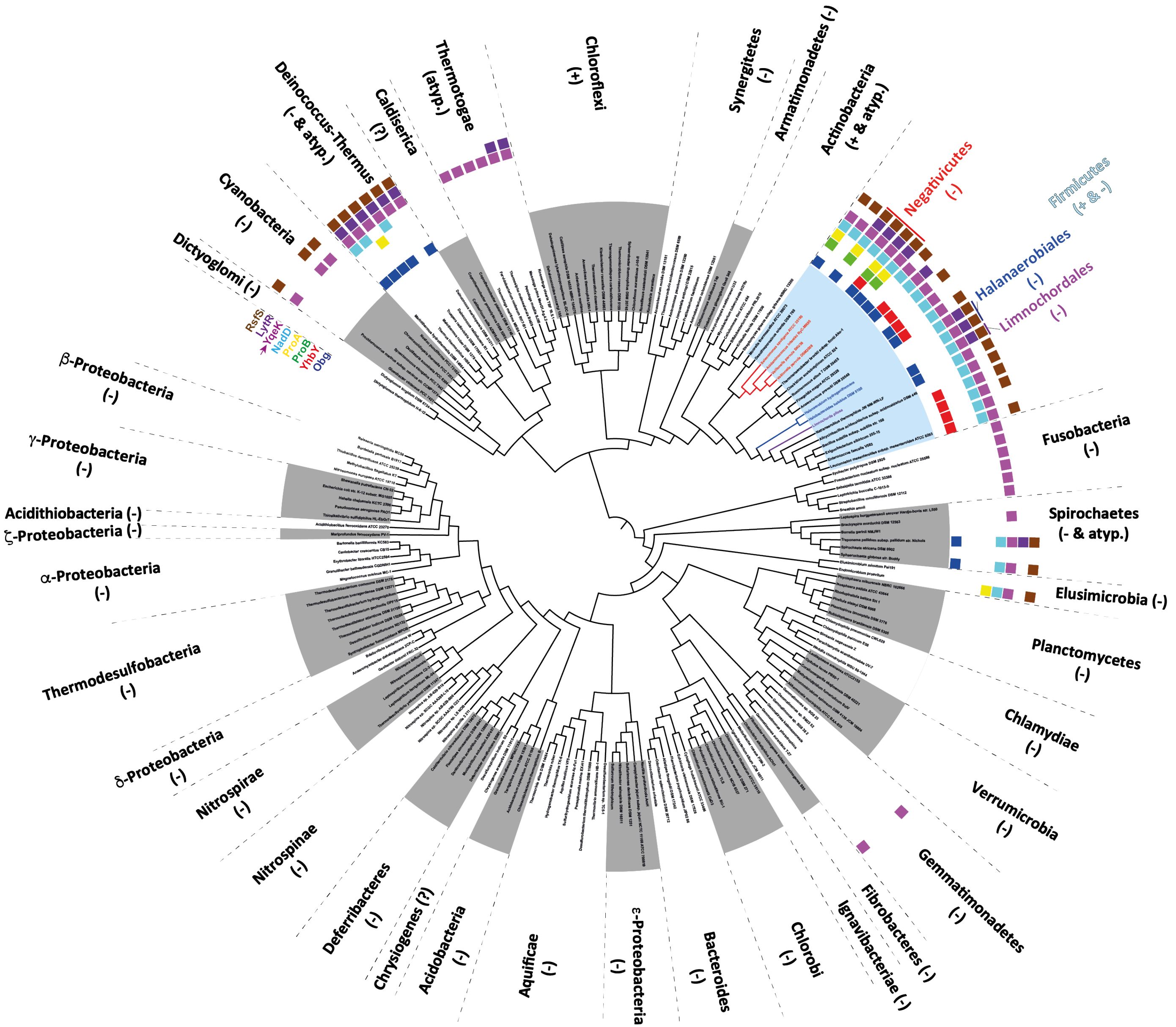

Chloroflexi
(+)
Synergistetes
(-)
Armatimonadetes (-)
Actinobacteria
(+ & atyp.)
Negativicutes
(-)
Firmicutes
(+ & -)
Halanaerobiales
(-)
Limnochordales
(-)
Fusobacteria
(-)
Spirochaetes
(- & atyp.)
Elusimicrobia (-)
Planctomycetes
(-)
Chlamydiae
(-)
Verrumicrobia
(-)
Gemmatimonadetes
(-)
Fibrobacteres (-)
Ignavibacteriae (-)
Chlorobi
(-)
Bacteroides
(-)
ε-Proteobacteria
(-)
Aquificae
(-)
Acidobacteria
(-)
Chrysiogenes (?)
Deferribacteres
(-)
Nitrospinae
(-)
Nitrospirae
(-)
δ-Proteobacteria
(-)
Thermodesulfobacteria
(-)
α-Proteobacteria
(-)
ζ-Proteobacteria (-)
Acidithiobacteria (-)
γ-Proteobacteria
(-)
β-Proteobacteria
(-)
RsfS
LytR
YqeK
NadD
ProA
ProB
YhbY
Obg
Dictyoglomi (-)
Cyanobacteria
(-)
Deinococcus-Thermus
(- & atyp.)
Caldiserica
(?)
Thermotogae
(atyp.)

Figure S5**: HD Phosphatase (YqeK) occurrence and synteny in diderm and monoderm bacteria**. **A.** The presence of the cluster was investigated using MacSyFinder (7) and the results were plotted onto a schematic tree of bacteria. A local databank containing 390 genomes representative of bacterial diversity was mined for the presence of a phosphatase containing HD domain (PF01966) using HMMSEARCH and the --cut_ga option. Protein sequences were then filtered using alignment, functional annotation, protein domains presence and phylogeny. Synteny was investigated in the locus around *yqeK* by looking, using MacSyFinder (7), for the presence of at least one of the 7 genes surrounding *yqeK* in *V parvula* SKV38, namely *obg* (containing GTP1_OBG domain, PF01018), *yhbY* (containing CRS1_YhbY domain, PF01985), *proB* (containing AA_kinase domain, PF00696), *proA* (containing Aldedh domain, PF00171), *nadD* (containing CTP_transf_like domain, PF01467), *lytR* (containing LytR_cpsA_psr domain, PF03816) and *rsfS* (containing RsfS domain, PF02410), with no more than eight other genes separating them. All HMM profiles were downloaded from the PFAM site (pfam.xfam.org). The results are plotted onto a schematic bacterial tree of 187 cultivable bacteria among the 390 of the analyzed databank. The cell wall status of each phylum is indicated as: (-) diderm with LPS, (+) monoderm, (atyp.) diderm without LPS, (?) unclear. For the Firmicutes, the diderm lineages are indicated in red (Negativicutes), blue (Halanaerobiales) and purple (Limnochordales).

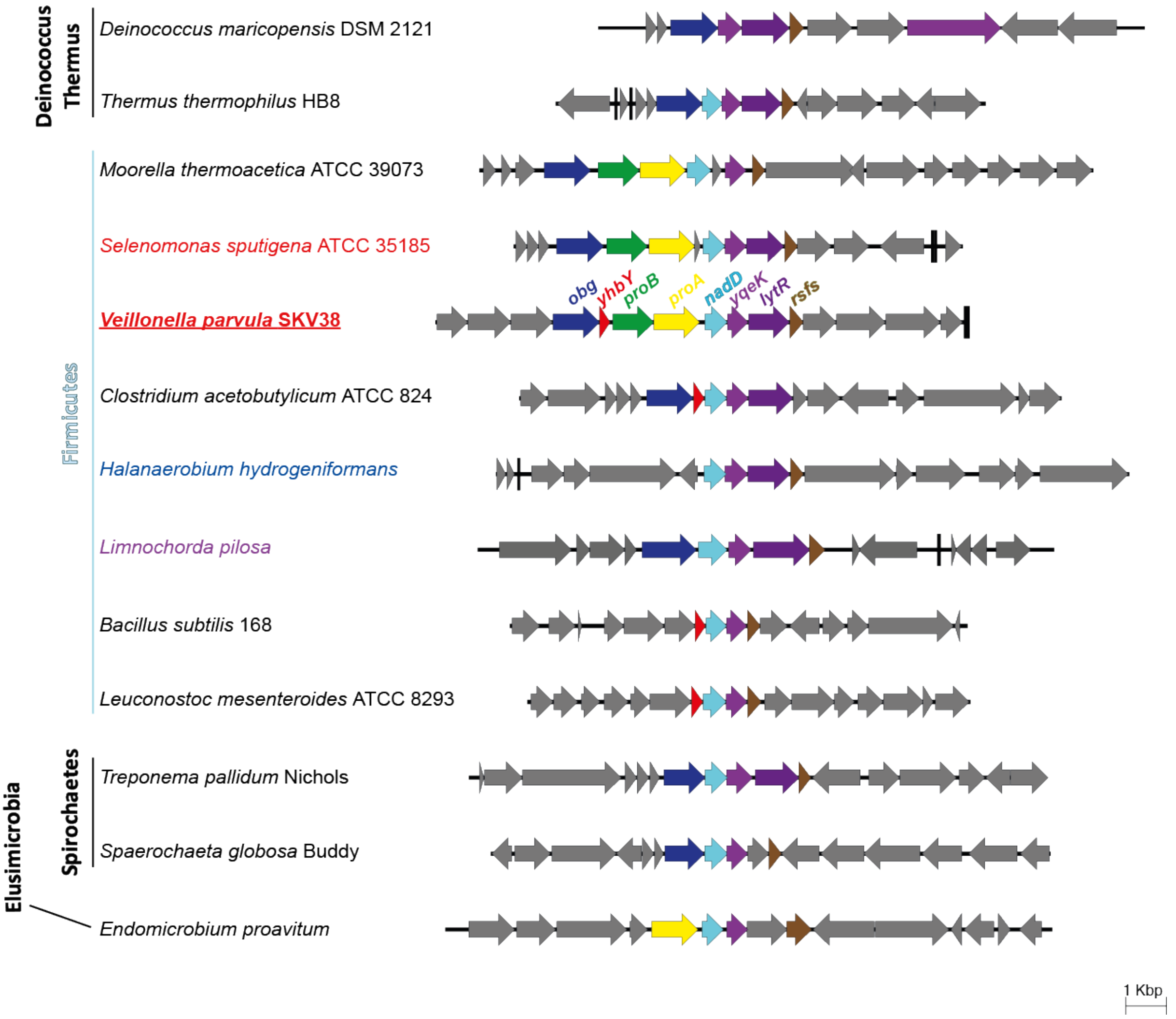


Figure S6**: Conservation of the HD Phosphatase encoding gene (*yqeK*) synteny in selected diderm and monoderm bacteria**. Some species from Figure S5 that contain a minimum of three genes in addition to *yqeK* within the cluster were selected and the corresponding genetic organizations were represented according to their phylogenetic proximity. The different genes are represented with different colors. The color code is the same as that of Figure S5. *V. parvula* SKV38 strain is underlined. In this strain *yqeK* corresponds to *FNLLGLLA_01127*.
